# Pyrophosphate-containing Calcium Phosphates Negatively Impact Heterotopic Bone Quality

**DOI:** 10.1101/2024.12.16.628729

**Authors:** Martina Jolic, Isabella Åberg, Omar Omar, Håkan Engqvist, Thomas Engstrand, Anders Palmquist, Peter Thomsen, Furqan A. Shah

## Abstract

In bone, critical size defects pose substantial challenge in maxillofacial and orthopaedic reconstructions as they are incapable of spontaneous regeneration. Autologous bone grafts remain the “gold standard” in bone repair but their limitations and associated morbidities drive the development of bone graft substitute materials. Assessing the osteoinductive properties of these substitutes can help predict their success in bone regeneration applications. Calcium phosphates (CaP) are widely used bone graft substitutes due to their biocompatibility, osteoconductive properties, and potential for osteoinductivity. CaP materials containing monetite, beta-tricalcium phosphate (β-TCP), and a small amount of calcium pyrophosphate (Ca-PP) possess both osteoconductive and osteoinductive properties. However, the role of Ca-PP in osteoinduction and material degradation remains unexplored. This study investigates heterotopic bone formation in response to five CaP material compositions, maintaining a constant monetite and β-TCP ratio, with varying amounts of Ca-PP (0 – 12.5%). Twelve adult female sheep (*Ovis aries*) were subcutaneously implanted with constructs made of six CaP tiles interconnected by a Ti6Al4V frame and a control implant. Histological analysis, backscattered electron scanning electron microscopy, and Raman spectroscopy of samples retrieved at 12- and 52 weeks reveal that Ca-PP does not hinder heterotopic bone formation and minimally impacts CaP degradation. While monetite and β-TCP transform into apatite, the Ca-PP phase remains unchanged. The addition of Ca-PP to the CaP material influences heterotopic bone quality and inflammatory response during tissue regeneration.

## 1. Introduction

Bone possesses a remarkable capacity for healing and regeneration. However, certain conditions such as injury, tumour resection, or debridement of infected tissue can result in critical size defects incapable of spontaneous regeneration (1). In such cases, reconstruction at the site of injury relies on natural bone grafts (autologous and allogenic) or synthetic bone graft substitutes (e.g., calcium sulphates, calcium phosphates, bioactive glasses, etc.) (2). Currently, autologous bone grafts are considered the “gold standard” in bone repair and reconstruction, meeting both the mechanical and biological requisites of an ideal replacement material. However, their limited availability, high resorption, and high infection rates, as well as associated morbidities, highlight the importance of developing synthetic bone graft substitutes (3–6).

Bone has a complex hierarchical structure composed of organic (predominantly collagen type I), inorganic (carbonated apatite, CHAp), and cellular components (7). Bone mineral, a poorly crystalline apatite susceptible to ion substitutions and rich in carbonate, and type I collagen collectively form the fundamental unit of bone matrix – the mineralised collagen fibril (8, 9). These fibrils organise into larger fibre bundles, forming plywood-like sheets (lamellae) that surround the Haversian canal and, finally, assemble into the basic structural unit of bone – the osteon (10). The resulting hierarchical organisation gives bone its overall mechanical properties, providing a unique combination of strength and toughness that is difficult to replicate in synthetic materials. Among bone graft substitutes, calcium phosphate (CaP) materials are extensively used due to their similarity to bone mineral, biocompatibility, biodegradability, and for their osteoconductive, and potential osteoinductive properties (11).

The concept of osteoinduction was first described by Urist in 1965, who demonstrated bone formation upon implantation of demineralised bone matrix in the muscles of mice, rats, rabbits, and guinea pigs (12). Consequently, an osteoinductive material was defined as a material supporting the osteogenic differentiation of progenitor cells (13). Currently, intrinsically osteoinductive materials are defined by their ability to mineralise *in vivo*, the presence of pores that can accommodate vascularisation, and their ability to create a local environment depleted of Ca^2+^ and PO_4_^3-^ (14). For a material to be considered osteoinductive, it should be able to induce bone formation heterotopically (15). While all osteoinductive materials are osteoconductive, the reverse is not always true. Rather, osteoconductive materials serve as effective scaffolds for new bone growth, with their effectiveness depending on the chemical composition and presence of macro- and microporosities (16). This distinction is especially relevant when working with CaP materials, as their bone regeneration capacity heavily relies on their osteoconductive as well as osteoinductive properties.

Various CaP compositions are already in clinical use, resulting in promising outcomes (17) and even outperforming autologous bone grafts in certain cases (18). Ongoing research into CaP materials aims to harness both their osteoconductive and osteoinductive properties to optimise bone regeneration strategies. Beta-tricalcium phosphate (β-TCP), recognized for its osteoconductive and osteoinductive properties, is one of the most commonly used bone graft substitutes in a wide variety of dental, orthopaedic, and drug delivery applications (19). Biphasic calcium phosphate (BCP) bioceramics, consisting of varying proportions of β-TCPs and hydroxyapatite (HAp), are among the most studied CaP-based bone graft substitutes (20), although the optimal β-TCP-to-HAp ratio remains debated. Higher β-TCP levels associated with faster biodegradation and higher levels of HAp impart better mechanical properties (21). The osteoinductive potential of CaP materials, particularly of BCPs with a HAp and β-TCP ratio of 60/40, was demonstrated in sheep models, with intramuscular bone formation observed after 6 months *in vivo* (22, 23). Additionally, monetite has shown favourable osteoconductive properties in various animal models as a component of mouldable cements, coatings, granules, and scaffolds (24). Another CaP phase of interest, calcium pyrophosphate (Ca-PP), has been shown to stimulate bone formation as a component of bone cements in the treatment of bone defect models (25). Porous Ca-PP scaffolds have displayed comparable osteoconductivity and improved *in vivo* degradation relative to HAp scaffolds in canine and rabbit models (26, 27). This is somewhat surprising, considering that ions of pyrophosphate are known inhibitors of mineralisation and mineral crystal growth, and the formation of Ca-PP crystals in articular cartilage is associated with inflammation, chondrocyte catabolism, and articular damage (28, 29). Moreover, the biological performance of CaP scaffolds can be easily modified through the inclusion of different mineral phases, ion doping, adjustment of pore sizes, or incorporation of growth factors (30).

We previously demonstrated that a monetite-based CaP material containing β-TCP and Ca-PP promotes bone regeneration in large cranial defects and guides bone formation beyond skeletal margins (31, 32). Furthermore, we established the osteoinductive properties of the material with similar composition at 12 weeks *in vivo* (31). Here, building on these findings, we investigated the osteoinductive capacity of a series of monetite-based CaP compositions with varying amounts of added Ca-PP over 12- and 52-week periods *in vivo*. Through detailed histological analysis and extensive Raman spectroscopy, we characterised the effects of Ca-PP on heterotopic bone formation and CaP material degradation.

## 2. Results

### 2.1. Quantitative histology

Undecalcified, Van Gieson-stained histological sections were used to determine the presence of heterotopic bone and measure the areas occupied by both heterotopic bone and CaP material constructs in samples retrieved after 12- and 52 weeks *in vivo*. Heterotopic bone was consistently found in samples, irrespective of the amount of Ca-PP in the material or the time point investigated (**Figure 1A-B**). In contrast, no heterotopic bone formed in response to the Ti6Al4V ELI control (**Supplementary Figures 12-13**). The exact values of heterotopic bone area (B.Ar.) and calcium phosphate area (CaP.Ar.) measured in sections of all compositions are detailed in **Supplementary Tables 2-3**. The measured B.Ar. was comparable between the various CaP material compositions at the respective time points, occupying an average area of 0.86 ± 0.77 mm^2^ and 1.63 ± 1.19 mm^2^, at 12- and 52 weeks *in vivo*, respectively (**Figure 1B**). Although there was a trend towards higher B.Ar. for all compositions at 52 weeks *in vivo* (p = 0.002), this difference was not statistically significant when comparing the two time points for individual CaP compositions. Nevertheless, substantial physical degradation of the CaP material occurred *in vivo* (**Figure 1C**). On average, the measured CaP.Ar. was 47.09 ± 13.08 mm^2^ at 12 weeks and was significantly reduced (p < 0.0001) to 12.75 ± 11.93 mm^2^ by 52 weeks.

**Figure 1.**
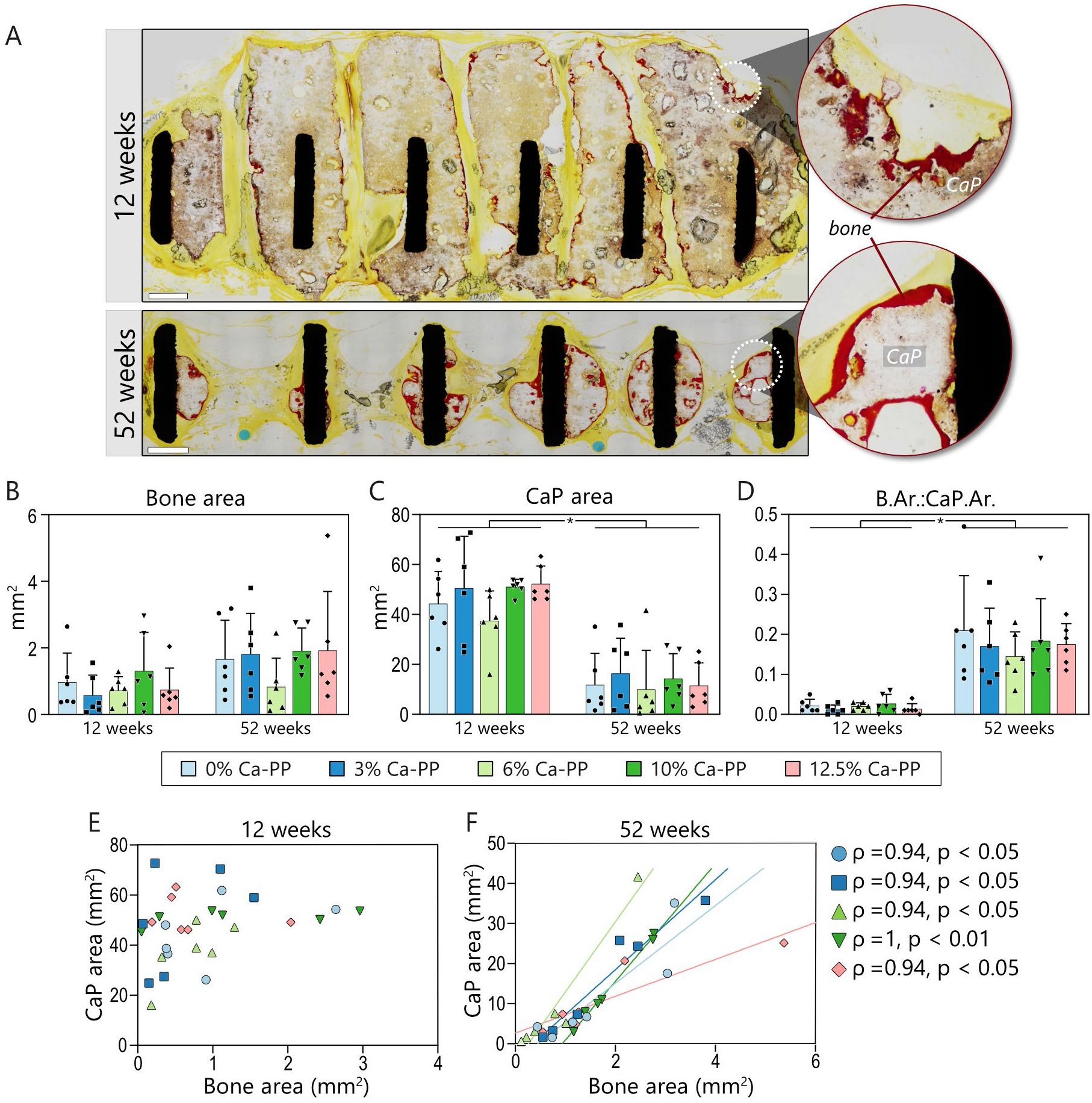
Heterotopic bone and CaP area in vivo. (A) Overviews of representative undecalcified histological sections stained with Van Gieson solution at 12- and 52 weeks (0% Ca-PP). Bone is stained intensely red, making it easily distinguishable from non-mineralised soft tissue and CaP. Scale bar = 1 mm. Histomorphometry: (B) heterotopic bone area (B.Ar.), (C) CaP area (CaP.Ar.), and (D) B.Ar. normalised to CaP.Ar. for respective CaP material compositions. Statistically significant difference marked with an asterisk, p = 0.002. (E) Relationships between CaP.Ar. and B.Ar. measured in respective CaP compositions at 12 weeks. (F) Relationships between CaP.Ar. and B.Ar. measured in respective CaP compositions at 52 weeks. Spearman’s rho correlation coefficient analysis.

Normalising B.Ar. to CaP.Ar. highlighted the time-dependent increase in heterotopic B.Ar., as shown in **Figure 1D**. At both 12- and 52 weeks, the measured CaP.Ar. was comparable between the CaP material compositions, irrespective of the Ca-PP content. While a positive relationship was not evident between B.Ar. and CaP.Ar. at 12 weeks *in vivo* (**Figure 1E**), at 52 weeks, the presence and the extent of heterotopic B.Ar. correlated with the remaining CaP material *in vivo*. Spearman’s correlation analysis confirmed this finding for all compositions (**Figure 1F**). Given that the amount of heterotopic bone formation was not influenced by the amount of Ca-PP in the CaP material, data from all groups were combined in further analyses. Pooled data revealed a low correlation between B.Ar. and CaP.Ar. at 12 weeks (r = 0.41, p < 0.05) that increased to a very high correlation at 52 weeks (r = 0.92, p < 0.0001).

### 2.2. Fate of CaP at a distance from the bone-material interface

Significant physical degradation of CaP from 12-to 52 weeks *in vivo* appears to be independent of the amount of Ca-PP originally present in the CaP material constructs. To determine the expected chemical transformation of the CaP material, point measurements were made in areas of the material located farthest from the bone-material interface using Raman spectroscopy (**Figure 2**). After 12 weeks *in vivo*, distinctive spectral features representing the three constituent phases, i.e., monetite (900 and 986 cm^−1^ peaks), β-TCP (947 and 970 cm^−1^ peaks), and Ca-PP (732 and 1043 cm^−1^ peaks), were identified. The intensity of Ca-PP peaks in the Raman spectra reflected the amount of Ca-PP present (0–12.5%) in the initial CaP material compositions. At 12 weeks, a partial transformation of the material to apatite was observed, as indicated by the inclusion of the ∼960 cm-1 (𝒱_1_PO_4_^3-^) peak in the Raman spectra. The spectra of the respective compositions were largely comparable between the two time points (**Figure 2**). However, at 52 weeks *in vivo*, a more pronounced chemical transformation of CaP was noted in the 0% and 6% Ca-PP compositions. In these, carbonated apatite (CHAp) was formed via the depletion of the monetite phase, as evidenced by the absence of peaks at 900 and 986 cm^−1^ and the emergence of the peak around ∼1070 cm-1 (𝒱_1_CO_3_^2-^). Conversely, the β-TCP and Ca-PP phases were still detectable within the averaged spectra (**Figure 2**).

**Figure 2.**
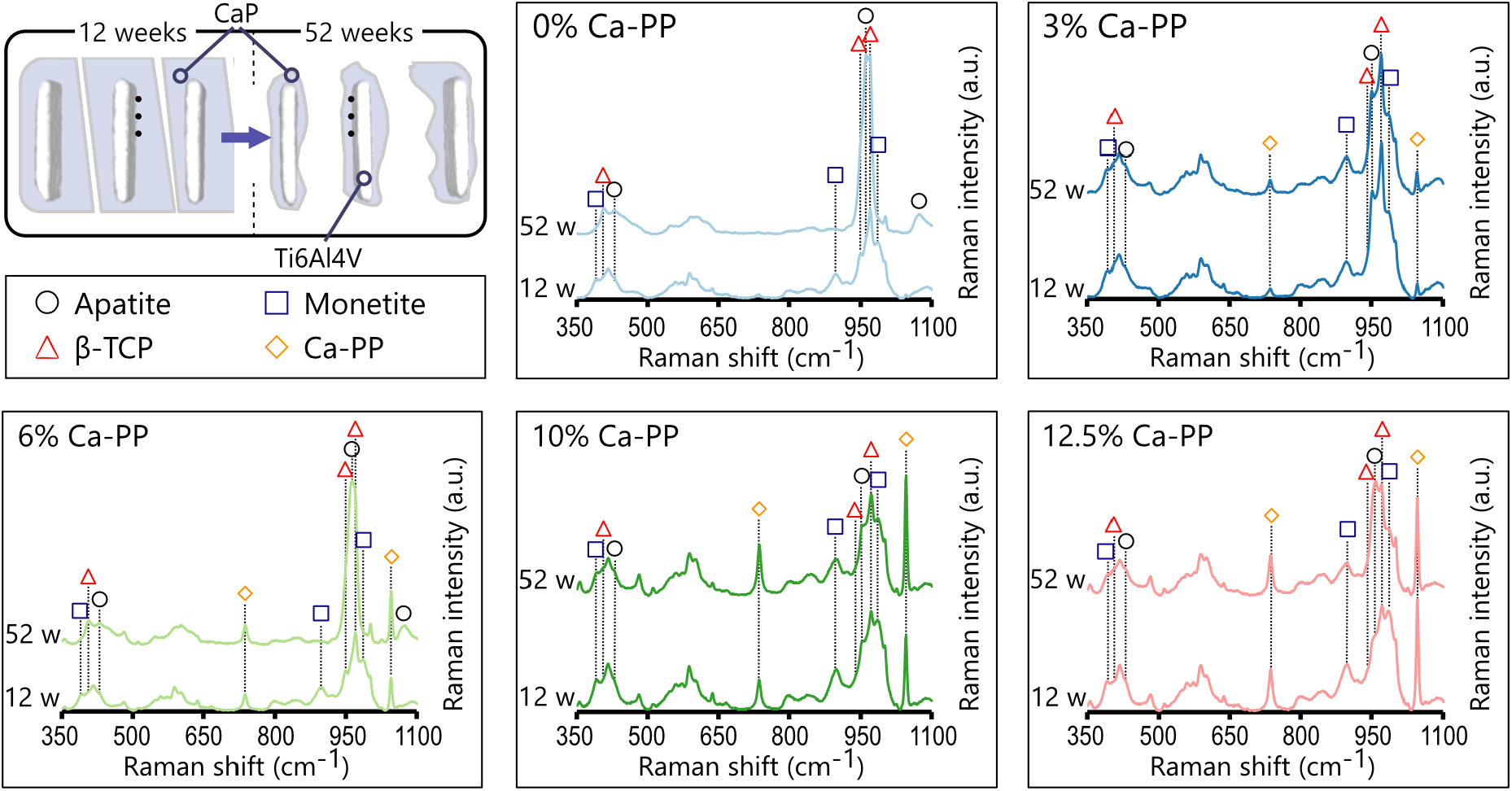
Fate of CaP in vivo. Top left: Schematic representation of significant CaP degradation *in vivo*. Black dots are representative of Raman spectroscopy point measurements in areas of the CaP material located farthest from the bone-material interface. Averaged Raman spectra (*n* = 6) of CaP at 12- and 52 weeks. Apatite (∼431 and ∼960 cm^−1^), carbonate (∼1070 cm^−1^), monetite (∼391, 900, and 986 cm^1^), β-TCP (∼407, 947, and 970 cm^−1^), and Ca-PP (∼732 and 1043 cm^−1^).

### 2.3. Fate of the CaP at the bone-material interface

The second region of interest (ROI) assessed with Raman spectroscopy was the CaP material in close proximity to the bone interface. At both 12- and 52 weeks *in vivo*, the CaP material at the interface underwent a substantial transformation into CHAp independent of the initial amount of Ca-PP (**Figure 3**). This likely occurred via the transformation of the monetite and β-TCP phases, as they were no longer detectable in the Raman spectra (**Figure 4**). At 52 weeks, from the original components of our CaP material, only the Ca-PP phase remained.

**Figure 3.**
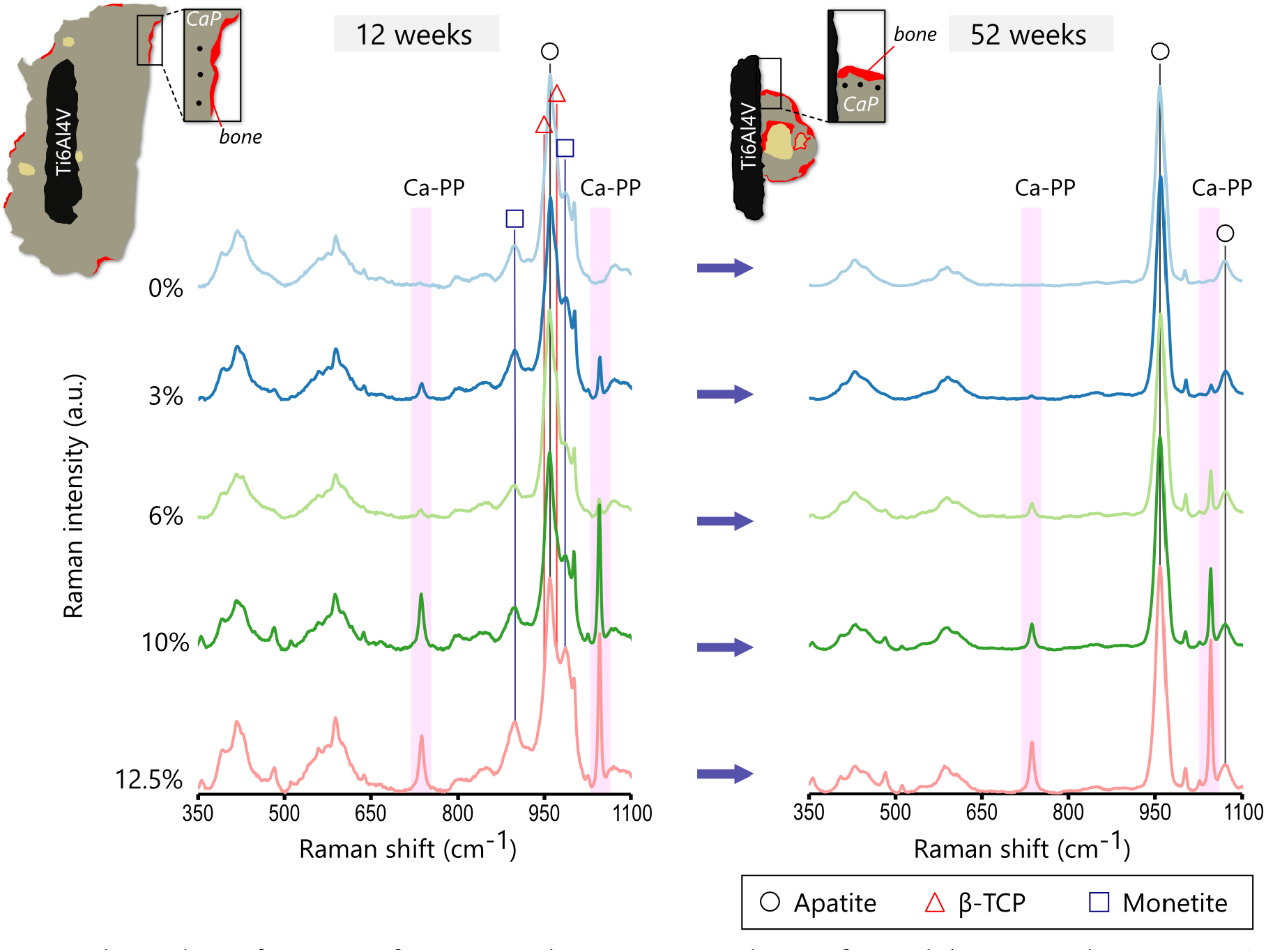
Chemical transformation of CaP material components at the interface with heterotopic bone. Top: Schematic representation of CaP material constructs at 12- and 52 weeks *in vivo* and the location of Raman spectroscopy point measurements (black dots) made in areas of the CaP material in close proximity (50 – 100 µm) to the bone interface. Averaged Raman spectra (*n* = 6) of CaP at 12- and 52 weeks. Apatite (∼960 cm^−1^), carbonate (∼1070 cm^−1^), monetite (∼900 and 986 cm^−1^), β-TCP (∼947 and 970 cm^−1^), and Ca-PP (∼732 and 1043 cm^−1^).

**Figure 4.**
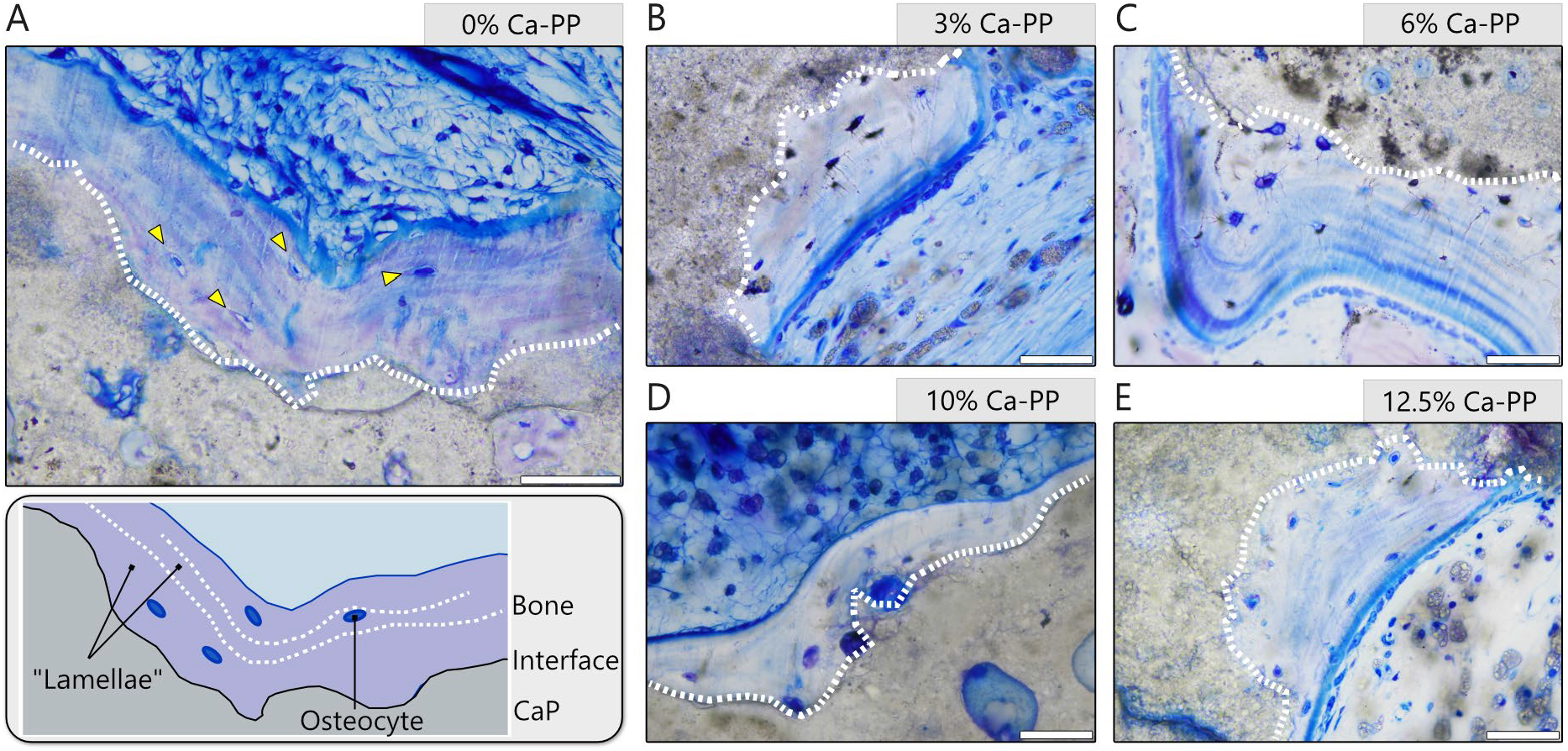
The lamellar-like appearance of heterotopic bone at 12 weeks in vivo. Representative, undecalcified, toluidine blue stained histological sections for: (A) 0% Ca-PP, (B) 3 % Ca-PP, (C) 6% Ca-PP, (D) 10% Ca-PP, and (E) 12.5 % Ca-PP. The interface between the heterotopic bone and the CaP material – white dotted line. Orientation of the osteocytes – yellow arrowheads. Scale bars = 50 µm.

### 2.4. Qualitative histology of heterotopic bone

Irrespective of the composition of the CaP material construct, at 12 weeks *in vivo*, a thin layer of heterotopic bone was found covering the material surface (**Figure 1A**). Toluidine blue staining of undecalcified histological sections revealed that at 12 weeks, bone followed the contours of the CaP material that had undergone partial degradation (**Figure 4**). In all compositions, the osteocytes were aligned parallel to the bone-material interface, and with the banded stain of the extracellular matrix, the lamellar-like appearance of bone was clearly evident (**Figure 4A-E**).

Osteoid, bone matrix that has not undergone the mineralisation process, was identified in all compositions at 12- and 52 weeks (**Figure 5**). At the surface of this newly formed bone matrix, osteoblasts were often observed, confirming ongoing bone formation. While osteoid areas were found at both time points, the incidence of these areas was reduced at 52 weeks.

**Figure 5.**
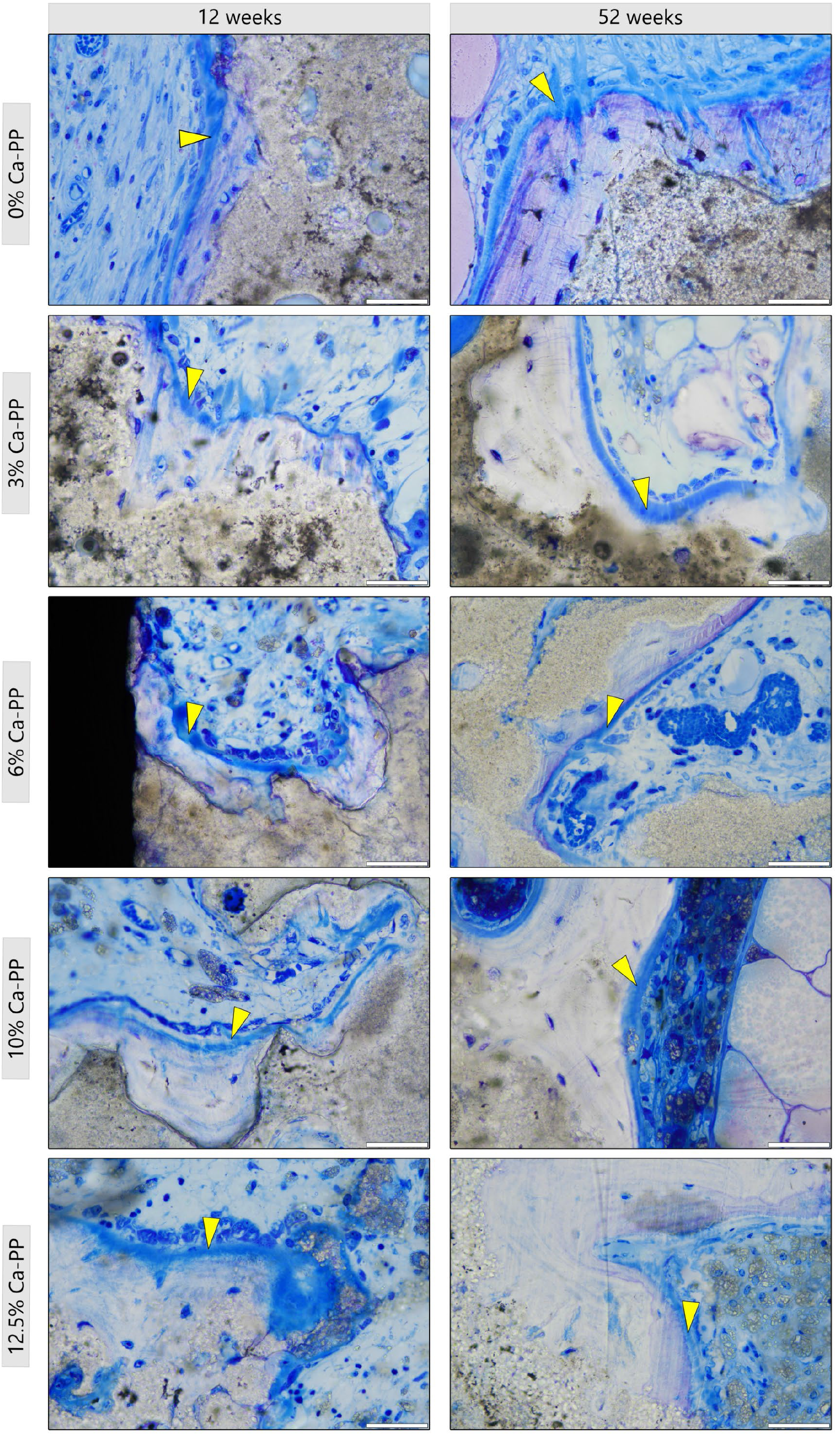
Presence of osteoid at 12- and 52 weeks in vivo. Representative, undecalcified, toluidine blue stained histological sections showing non-mineralised bone matrix, osteoid (yellow arrowheads), found with all CaP material compositions. Scale bars = 50 µm.

At 52 weeks *in vivo*, larger islands of bone were found. Moreover, large fat cells, adipocytes, were found in all investigated compositions (**Figure 6A-E**). These cells were observed not only in the soft tissue of the inter-tile space but also in large islands of bone within the remaining CaP material bulk. Seemingly, these adipocytes occupied areas in which heterotopic bone has undergone remodelling. Here, the lamellar bone structure was more obvious, while the material interface was more clearly demarcated.

**Figure 6.**
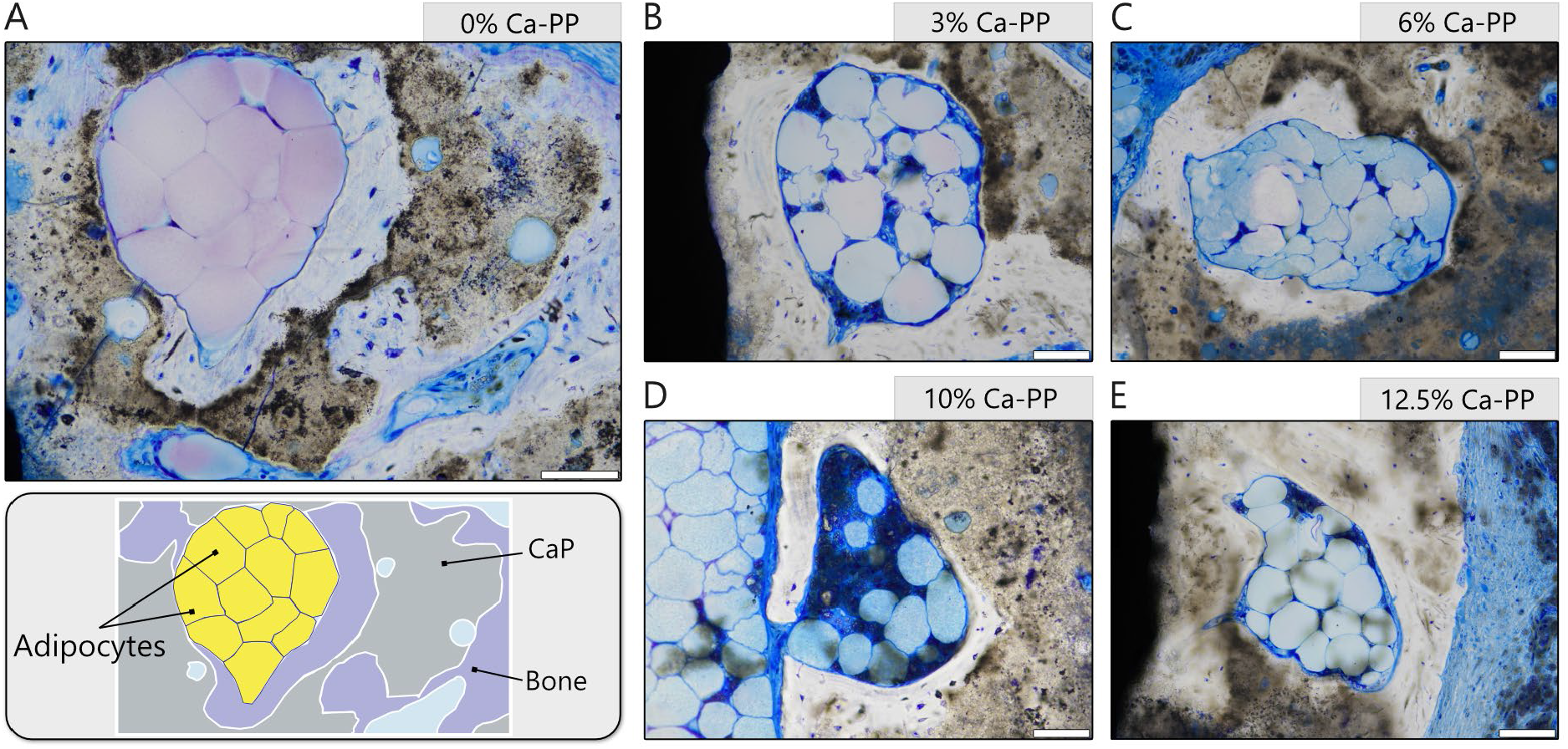
Presence of adipose cells at 52 weeks in vivo. Representative, undecalcified, toluidine blue stained histological sections showing the presence of the adipocytes in larger islands of heterotopic bone for: (A) 0% Ca-PP, (B) 3% Ca-PP, (C) 6% Ca-PP, (D) 10% Ca-PP, and (E) 12.5% Ca-PP at 52 weeks. Scale bars = 100 µm.

The CaP material underwent significant degradation *in vivo* (see **Figure 1C**), which appeared to be dependent not only on the exposure of the material to the surrounding environment (i.e., interstitial fluid) but also facilitated by cells. Multinucleated cells were often observed at the surface of the CaP material, with CaP particles clearly visible intracellularly at 12- and 52 weeks (**Figure 7, Supplementary Figure 14**). At 52 weeks, few multinucleated cells were observed in association with CaP constructs containing 0% Ca-PP, and in all other compositions, their numbers were reduced compared with 12 weeks *in vivo*.

**Figure 7.**
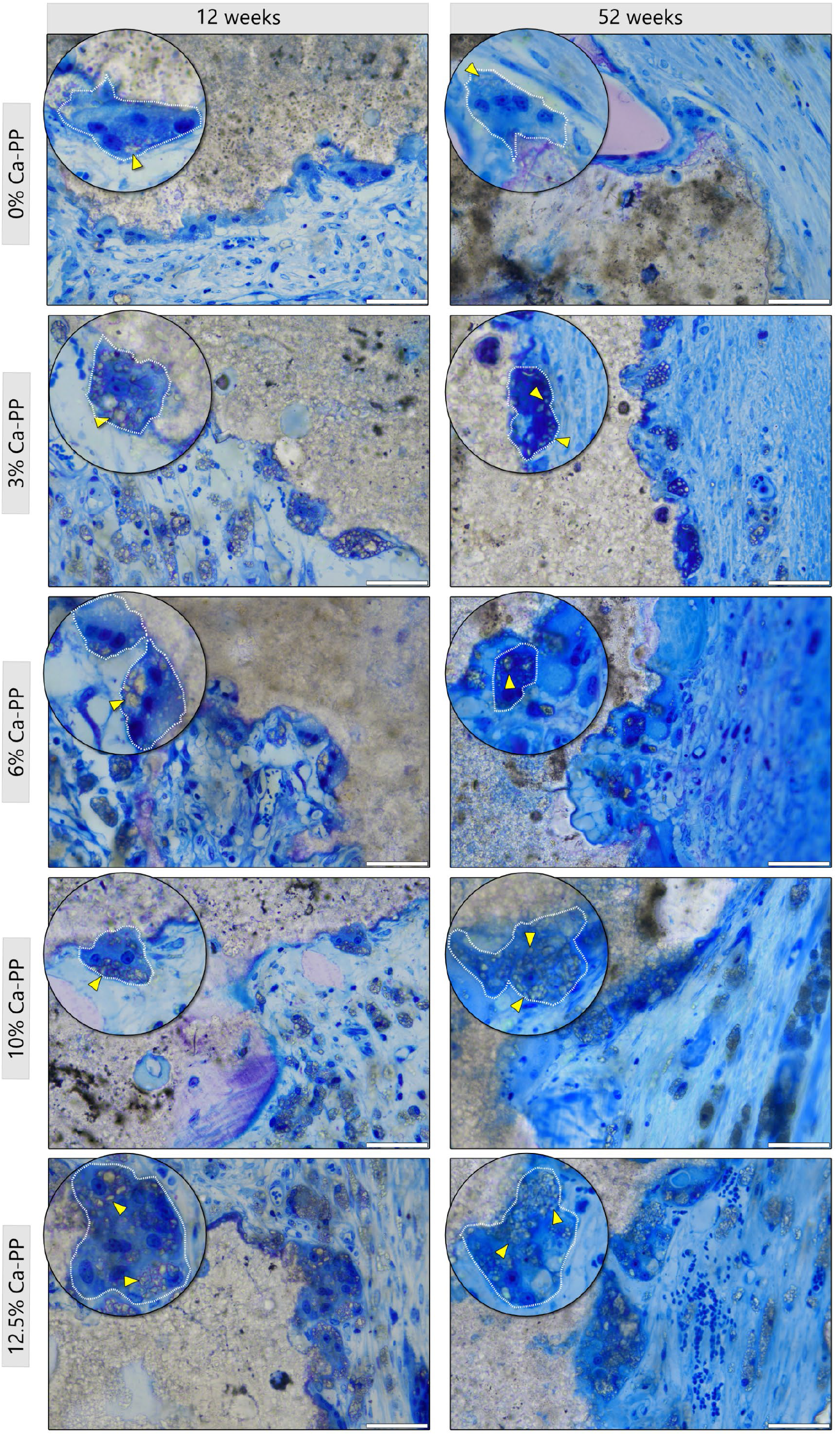
Calcium phosphate is actively removed at 12- and 52 weeks in vivo. Representative, undecalcified, toluidine blue stained histological sections showing multinucleated cells found near all CaP material compositions. Large particles of CaP material (yellow arrowheads) within the intracellular space. Scale bars = 50 µm.

### 2.5. Heterotopic bone characterisation

The chemical composition of heterotopic bone was investigated using Raman spectroscopy. At both 12- and 52 weeks, characteristic spectral features of the mineral (i.e., CHAp) and organic phases (i.e., amide III, proline, hydroxyproline) of bone were identified. The Raman spectra were comparable across the various CaP material compositions at 12- and 52 weeks (**Figure 8A**). Notably, we observed the incorporation of the Ca-PP peak in the Raman spectra of heterotopic bone, with the appearance of the ∼1043 cm^−1^ peak at both 12- and 52 weeks (**Figure 8A**).

**Figure 8.**
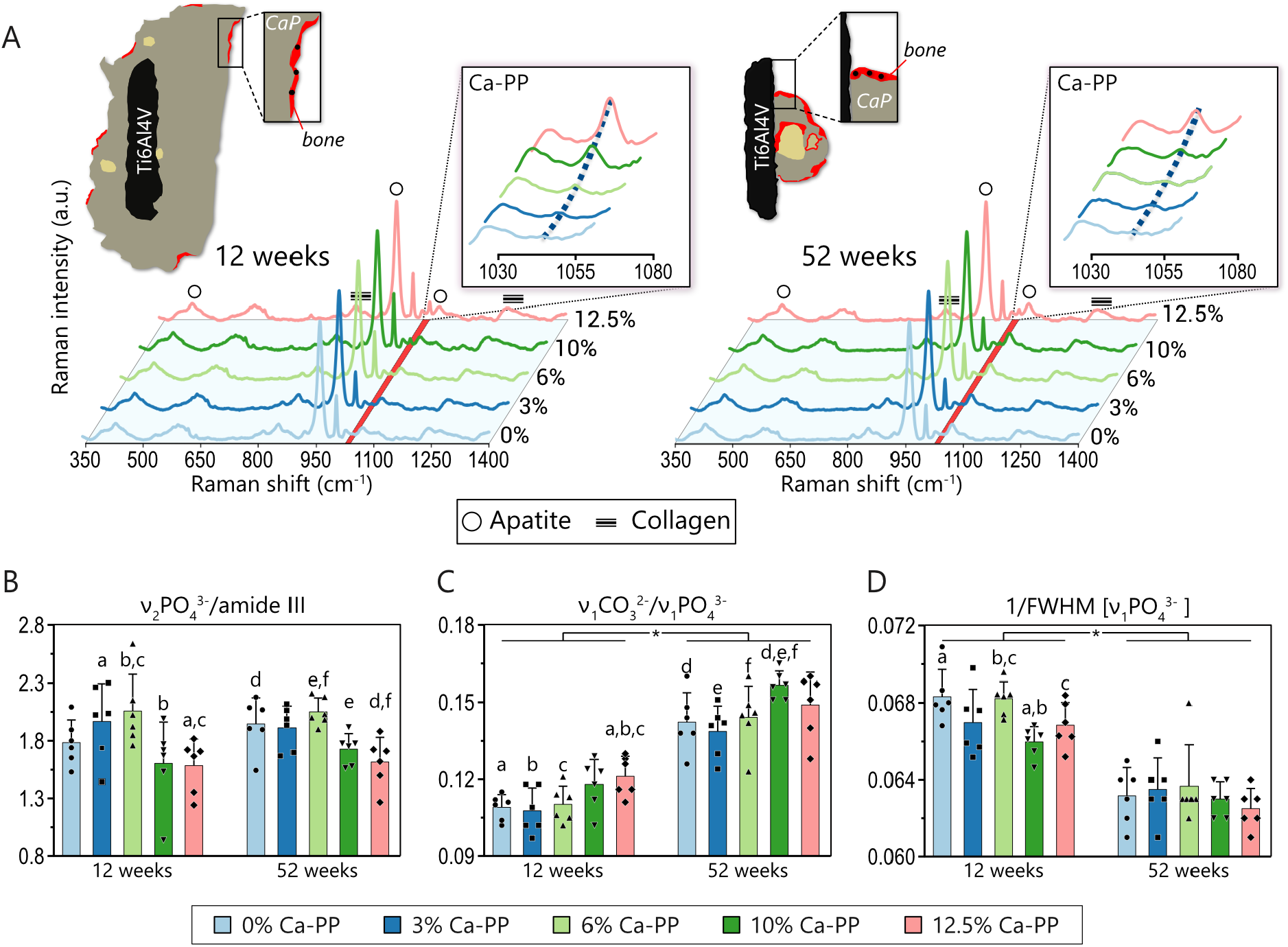
Chemical composition of heterotopic bone. (A) Top: Schematic representation of CaP material constructs at 12- and 52 weeks *in vivo* and locations of Raman spectroscopy point measurements (black dots) made in heterotopic bone at 12- and 52 weeks. Averaged Raman spectra (*n* = 6) for respective compositions (0–12.5% Ca-PP) were comparable at both time points. Characteristic spectral features of bone: apatite (∼960 cm-1), 𝓋_1_CO_3_ ^2-^ (∼1070 cm-1), amide III (∼1240–1270 cm-1), proline and hydroxyproline (∼850–873 cm^−1^). Insets show the 1030–1080 cm^−1^ range of Raman spectra and the appearance of the Ca-PP peak at 1043 cm^−1^. Extracellular matrix characterisation: (B) mineral-to-matrix ratio as the integral area ratio of 𝓋_2_PO43-(420–470 cm-1) and amide III band (1240–1270 cm-1), (C) carbonate-to-phosphate ratio as intensity ratio of 𝓋_1_CO32- (∼1070 cm-1) and 𝓋_1_PO43- (∼960 cm-1) peaks, and (D) mineral crystallinity as the inverse value of full-width-at-half maximum (FWHM) of the 𝓋_1_PO43- (∼960 cm-1) peak. The same letters indicate a significant difference between the two compositions. The asterisk indicates a significant difference between respective compositions at two time points.

The degree of mineralisation, as indicated by the mineral-to-matrix ratio, was mostly comparable between the CaP material compositions, with an average 𝓋_2_PO_4_^3-^/amide III ratio of 1.8 ± 0.3 and 1.85 ± 0.2 at 12- and 52 weeks, respectively. Generally, the degree of mineralisation was higher in response to constructs containing lower amounts of Ca-PP (0, 3, and 6%) (**Table 1**).

**Table 1.** Raman metrics tested in heterotopic bone at 12- and 52-weeks in vivo.

| Group | 12 weeks |  |  | 52 weeks |  |  |
| --- | --- | --- | --- | --- | --- | --- |
| | $\nu_2\text{PO}_4^{3-}$/<br>amide III | $\nu_1\text{CO}_3^{2-}$/<br>$\nu_1\text{PO}_4^{3-}$ | 1/FWHM<br>($\nu_1\text{PO}_4^{3-}$) | $\nu_2\text{PO}_4^{3-}$/<br>amide III | $\nu_1\text{CO}_3^{2-}$/<br>$\nu_1\text{PO}_4^{3-}$ | 1/FWHM<br>($\nu_1\text{PO}_4^{3-}$) |
| **0% Ca-PP** | $1.79 \pm 0.20$ | $0.11 \pm 0.01$ | $0.068 \pm 0.001$ | $1.95 \pm 0.23$ | $0.14 \pm 0.01$ | $0.063 \pm 0.001$ |
| **3% Ca-PP** | $1.97 \pm 0.33$ | $0.11 \pm 0.01$ | $0.067 \pm 0.002$ | $1.91 \pm 0.19$ | $0.14 \pm 0.01$ | $0.064 \pm 0.002$ |
| **6% Ca-PP** | $2.06 \pm 0.31$ | $0.11 \pm 0.01$ | $0.068 \pm 0.001$ | $2.05 \pm 0.12$ | $0.14 \pm 0.01$ | $0.064 \pm 0.002$ |
| **10% Ca-PP** | $1.60 \pm 0.36$ | $0.12 \pm 0.01$ | $0.066 \pm 0.001$ | $1.73 \pm 0.13$ | $0.16 \pm 0.01$ | $0.063 \pm 0.001$ |
| **12.5% Ca-PP** | $1.59 \pm 0.23$ | $0.12 \pm 0.01$ | $0.067 \pm 0.001$ | $1.62 \pm 0.21$ | $0.15 \pm 0.01$ | $0.063 \pm 0.001$ |

At 12 weeks, the mineral-to-matrix ratio was highest with 6% Ca-PP, with significant differences seen when compared with 10% and 12.5% Ca-PP (p < 0.05 and p < 0.01, respectively) (**Figure 8B**). The second highest ratio was seen with 3% Ca-PP (p < 0.05 vs 12.5% Ca-PP) (**Table 1**). At 52 weeks, the mineral-to-matrix ratio was also highest with 6% Ca-PP (p < 0.01 vs 10%- and 12.5% Ca-PP), while the second highest ratio was seen with 0% Ca-PP (p < 0.05 vs 12.5% Ca-PP). At both time points, the lowest mineral-to-matrix ratio was observed with 12.5% Ca-PP.

For all compositions, the carbonate-to-phosphate ratio significantly increased with time. This increase reflects a B-type carbonate substitution of PO_4_^3-^ for CO_3_^2-^ in the apatite structure, leading to changes in size and increased solubility of the mineral (8). We found significantly lower carbonate-to-phosphate ratios with constructs containing low amounts of Ca-PP at both time points (**Table 1**).

At 12 weeks, the carbonate-to-phosphate ratio with 12.5% Ca-PP was significantly higher than 0-, 3-, and 6% Ca-PP (p < 0.05). At 52 weeks, the highest carbonate-to-phosphate ratio was found with 10% Ca-PP, which was significantly higher than 0-, 3-, and 6% Ca-PP (p < 0.05 vs 0- and 6% Ca-PP, and p < 0.01 vs 3% Ca-PP) (**Figure 8C**).

The mineral crystallinity (1/FWHM) decreased from 12-to 52 weeks *in vivo* (**Table 1**). At 12 weeks, significant intergroup differences could be calculated. The mineral crystallinity was significantly higher with 6% Ca-PP than 10- and 12.5% Ca-PP (p < 0.01) and with 0% Ca-PP than 10% Ca-PP (p < 0.01) (**Figure 8D**).

## 3. Discussion

Large bone defects caused by trauma or tumour resections remain a significant problem in maxillofacial and orthopaedic surgery despite the significant advances made in the area of bone repair and reconstruction. Osteoinduction is the process of inducing new bone tissue formation, and a material is considered osteoinductive if it can stimulate bone formation in locations where bone does not typically exist, such as beneath the skin or within the muscle tissue. Testing materials for their osteoinductive capacity is essential for determining their efficacy and biocompatibility for bone augmentation, establishing their ability to integrate into the surrounding bone tissue and enhance normal healing processes, and predicting their clinical success. The heterotopic bone formation discussed here occurs in response to the presence of an osteoinductive material and should not be mistaken for heterotopic mineralisation, which includes ossification in pathological conditions (36).

In our study, we tested the osteoinductive properties of a multicomponent CaP material implanted in a subcutaneous sheep model. Different compositions of the material were prepared, with fixed amounts of monetite and β-TCP in a 10:1 ratio and varying amount of Ca-PP (0 – 12.5%). The combination of monetite and β-TCP is known to support heterotopic ossification (i.e., osteoinduction), on the other hand, inorganic pyrophosphate ions have long been recognised as inhibitors of apatite precipitation and mineralisation because of their role in preventing crystal nucleation (37, 38). We found that the incorporation of Ca-PP in the multicomponent CaP material had minimal impact on heterotopic bone formation. After 12- and 52 weeks *in vivo*, heterotopic bone was consistently detected across all compositions, irrespective of the amount of Ca-PP added, although the quality of *de novo* bone was compromised with higher amounts of Ca-PP. Moreover, we saw no heterotopic bone formation using control implants made of Ti6Al4V ELI at both time points, highlighting the osteoinductive properties of the tested multicomponent CaP material.

It has been reported that the addition of a high amount of Ca-PP (i.e., 28 wt.%) to brushite cements enhances bone formation in a sheep bone-defect model (25). In this example, poor interfacial adhesion between the Ca-PP particles and brushite probably lowers the durability of the material, which facilitates the creation and dispersal of material fragments. This process thereby increases the available surface area for further material degradation, favouring osteoinduction and a larger *de novo* bone area (25). Here, we did not observe a similar osteogenic effect of Ca-PP, as the amount of Ca-PP added (0 – 12.5%) did not influence the extent of CaP material degradation. This is further highlighted by a time-dependent increase in B.Ar. from 12-to 52 weeks *in vivo* that reached statical significance only upon normalisation to CaP.Ar. The increase in the normalised B.Ar. value was a result of the substantial CaP material degradation after 52 weeks *in vivo*, with a direct correlation observed between the B.Ar. and CaP.Ar.

It is generally assumed that when a bone repair biomaterial degrades, the resulting voids allow the in-growth of bone, whereby the amount of remaining implant material is inversely correlated with the area (or volume) occupied by newly formed bone. A characteristic feature of bone regeneration within bone defects using biomaterials is that while occupying an enclosed space, any newly formed bone is *attached* to both the implant material and the native bone. In contrast, without confinement by the native bone, such as in a soft tissue implantation site, the newly formed bone is primarily attached to the implant material. Upon degradation, fragments of the implant material are *cleared* (or removed) from the local site along with the newly formed attached bone. In other words, degradation of the implant material is expected to contribute to a partial loss of the newly formed bone. Therefore, the amount of heterotopic bone amenable to quantitative analysis is dependent on the amount of the remaining implant material.

One of the most sought-after properties of bone substitute materials is a degradation rate that matches that of new bone formation (39). Therefore, understanding the degradation process of the investigated material is of special importance. Following implantation, all implanted CaP material constructs underwent substantial physical degradation *in vivo*, the extent of which was unrelated to the Ca-PP content. At least two mechanisms contribute to this process: (*i*) passive material degradation driven by factors such as mechanical stresses from the physiological environment and (*ii*) active material degradation involving cells. Solubility is a key factor in the loss of mass of CaP materials with low porosity during *in vivo* degradation (40). Monetite, which constitutes ∼80–91% in this series of compositions, may be the main driving force behind the degradation of the CaP material between 12- and 52 weeks *in vivo*, given its greater solubility than that of both β-TCP and Ca-PP at neutral pH (41, 42).

On the surface and in close proximity to the implanted constructs, we observed multinucleated cells with aggregates of CaP fragments, which were easily discernible in the intracellular space, both at 12- and 52 weeks. In similar Ca-PP containing CaPs, this intracellular debris has been shown to consist exclusively of Ca-PP particles (32, 43). In our work, we did not find that the extent of material bulk degradation is changed by increasing the amount of Ca-PP, although higher amounts of Ca-PP appear to exacerbate the phagocytosis of the CaP material. We noted that the area occupied by multinucleated cells with intracellular CaP debris increased with the amount of added Ca-PP from 3 – 12.5 %, whereas with 0% Ca-PP, these cells were scarcely found. A similar observation was made in a subcutaneous rat model where higher amounts of phagocytosed microparticles were found with four β-TCP implants doped with increasing amounts of Ca-PP (0.5 – 10 wt%). A higher amount of Ca-PP was associated with the presence of inflammatory cells at 4 weeks following implantation (44). Here, we did not observe an increase in inflammation during visual inspection of the implantation site at sample retrieval, and in the histological investigations the increased presence of multinucleated cells was linked to material degradation process.

We also found that the CaP material underwent substantial chemical transformation *in vivo*. Depending on the pH and composition of the microenvironment, CaPs can transform from one phase to another (45). Here, after 12 weeks *in vivo*, Raman spectroscopy revealed the presence of apatite in addition to the constituent phases of the multicomponent CaP material. This transformation is attributed primarily to the dissolution of monetite, as β-TCP is insoluble in physiological conditions. However, the solubility of β-TCP increases at acidic pH, leaving cell-mediated resorption as a main mode of dissolution (19). After 52 weeks, a near-complete transformation was noted at the CaP material-bone interface, with apatite being the predominant CaP phase detected. However, the presence of Ca-PP peaks within the Raman spectra points toward low solubility of this phase *in vivo* and highlights the importance of phagocytosis as a primary means of Ca-PP removal in non-osseous sites.

In areas of the CaP material located farthest from the bone interface, we found that the transformation to apatite was less pronounced at 12 weeks *in vivo*, confirming that exposure to cellular activity increases the chemical transformation of the material. Nevertheless, at 52 weeks *in vivo*, the material had undergone considerable transformation at these distant sites. In the compositions with 0- and 6% Ca-PP, the monetite phase was no longer detectable in the averaged Raman spectra, even though it could be observed in the individual point measurements. In contrast, in the compositions with 3-, 10-, and 12.5% Ca-PP, the averaged Raman spectra showed the presence of all initial phases (i.e., monetite, β-TCP, and Ca-PP). As the selection of ROIs was performed using backscattered electron scanning electron microscopy (BSE-SEM) images, the obtained information is limited to component distribution (i.e., CaP, bone, and surrounding tissues) at the sample surface. It is possible that the greater extent of degradation observed in compositions with 0- and 6% Ca-PP compositions at 52 weeks was due to the sampled sites being in closer proximity to bone and/or interstitial fluid than intended, introducing an unintentional bias into our measurements.

Several questions regarding the long-term fate of the CaP material remain, specifically: (*i*) what happens with the phagocytosed CaP material during cell mitosis or apoptosis, (*ii*) will the observed multinucleated cells still be present in the subcutaneous pocket once the remaining bulk CaP material has been degraded, and (*iii*) what is the fate of the already phagocytosed CaP material if it cannot be resorbed? Research on phagocytosis of undegradable microparticles has shown that such particles can be divided between daughter cells during mitosis, and in case of apoptosis, released particles can be taken up by other macrophages (46). It remains to be seen if intracellular CaP debris can be resorbed with prolonged exposure to the acidic environment in the phagosome or if a similar scenario could apply here. It is expected that the ongoing degradation of the implanted CaP material will continuously provide new debris for phagocytosis. During degradation *in vivo*, BCPs release HA particles that are phagocytosed by macrophages but are not effectively degraded, leading to local tissue inflammation and cell damage (47). It stands that if phagocytosed CaP material particles are resistant to dissolution and degradation, they will likely contribute to cellular stress and apoptosis, triggering the release of the inflammation mediators, inducing repeated phagocytosis, and ultimately leading to fibrous encapsulation of the particles in the subcutaneous pocket.

Beyond the effect on degradation of the CaP material, we investigated the impact of Ca-PP on heterotopic bone formation and quality. At 12 weeks, heterotopic bone formation was restricted mainly to the outer regions of the implants and was rarely observed within the bulk of CaP material. This is most likely caused by the low interconnected porosity of our material. Here, the heterotopic bone formed through intramembranous ossification, with no evidence of intermediate cartilage tissue found at either time point. At 12 weeks *in vivo*, heterotopic bone followed the contours of the partially degraded CaP material and had a lamellar-like organisation. The lamellar-like appearance is interpreted as coordinated bone formation activity resulting from the initial organisation of osteoblasts on the surface of the substrate (e.g., CaP material), which plays a critical role in the formation of ordered tissues (48). The bone continuously formed on the material surface. In toluidine blue-stained histological sections, non-mineralised bone tissue (i.e., osteoid) was found across all compositions at both 12- and 52 weeks *in vivo*. In the majority of cases, osteoid was observed with associated polarised surface osteoblasts. However, the incidence of such observations was reduced at 52 weeks for all CaP material constructs regardless of the amount of Ca-PP. This sustained bone formation is attributed to the continuous degradation of the CaP material, including the development of cracks within the material, particle dispersal, and release of Ca^2+^ and P^−^.

After 52 weeks *in vivo*, large islands of bone were found in all compositions. This bone occupied both the surface concavities and large pores within the degraded CaP material. The most striking observation in all compositions was that of large adipocytes in bone areas that had undergone a remodelling process, creating compartments resembling bone marrow. It is well-documented that disuse osteopenia, i.e., bone loss due to local skeletal unloading, is associated with the accumulation of fat in the bone marrow (49, 50). We presume that the accumulation of adipocytes in heterotopic bone in our study is caused by the lack of mechanical loading within the subcutaneous pocket. During normal bone remodelling within the skeletal system, the amount of resorbed bone is in net balance with the amount of newly formed bone to ensure the structural integrity of the tissue (51). However, here, it appears that once the strong pro-osteogenic microenvironment generated by the material-induced immune response subsided, the remodelling processes that took place favoured the differentiation of mesenchymal cells into adipocytes. In another study, heterotopic bone, formed in response to HAp and BCP implants, and bone-like tissue in β-TCP implants, were found as long as 2.5 years *in vivo* after implantation in the dorsal muscles of dogs (52). Similar to our findings, the formed bone showed clear signs of remodelling and the presence of a bone marrow compartment. Both in that study and here, the heterotopically formed bone was found in the presence of implanted materials, confirming that although long-term survival of heterotopic bone is possible, it is contingent on the continued presence of the CaP material in a non-osseous site.

The presence of Ca-PP did not affect the composition of the heterotopic bone formed at either 12- or 52 weeks *in vivo*, with typical organic and mineral spectral features of extracellular bone matrix present in Raman spectra. However, we observed the presence of the Ca-PP peak (∼1043 cm^−1^) in the Raman spectra of the extracellular bone matrix. In a previous report, this Ca-PP peak was found in the Raman spectra of extraskeletal bone formed in response to CaP constructs with a similar composition, which was ascribed to the presence of discrete Ca-PP particles embedded within the bone matrix (32). Here, we noted a positive relationship between the intensity of the Ca-PP signal in the Raman spectra and the amount of Ca-PP in the initial CaP mixture, suggesting that higher initial amounts of Ca-PP lead to greater incorporation of discrete particles of Ca-PP within the heterotopic bone matrix. Still, our findings suggest that the inclusion of Ca-PP in an osteoinductive material may negatively affect the level of mineralisation of *de novo* bone (**Figure 8**). At both 12- and 52 weeks, mineralisation levels of heterotopic bone were the lowest in compositions with the highest amounts of Ca-PP (i.e. 10- and 12.5%). Although the mineral-to-matrix ratio, as well as the carbonate-to-phosphate ratio, generally increase with tissue age (53–55), here, the mineral content did not increase with time from 12-to 52 weeks. While this could be due to a process where CaP material is continuously degraded and *de novo* bone is consequently formed, the presence of larger islands of bone tissue with osteonal organisation and clear signs of remodelling in the heterotopic bone at 52 weeks render such a scenario unlikely. Both these observations suggest a more “mature” bone tissue at 52 weeks, contrasting with the lamellar-like bone on the surface of the constructs at 12 weeks. Alternative explanation for the comparable levels of mineralisation at the two time points is that amounts of mineral and collagen increased proportionally within the extracellular bone matrix of heterotopic bone. This notion is supported by the higher carbonate-to-phosphate ratio of bone found in all compositions at 52 weeks (p < 0.01). We observed a gradual increase in B-type carbonate substitution in the heterotopic bone from 0-to 12.5% Ca-PP, with the highest values measured with 12.5- and 10% Ca-PP at 12- and 52 weeks, respectively. It appears that higher amounts of Ca-PP within the extracellular bone matrix resulted in delayed bone formation and/or bone remodelling. Furthermore, the incorporation of CO_3_^2-^ into apatite lattice negatively correlated with mineral crystallinity (1/FWHM). Generally, substitutions in the apatite lattice structure are indicative of changes in the perfection and size of crystallites and are negatively correlated with mineral crystallinity(56, 57). However, large changes in the CO_3_^2-^ content do not always translate to large impacts on the FWHM of 𝓋_1_ PO_4_^3−^ (58). At 12 weeks, the lowest mineral crystallinity was observed in the bone that formed in response to constructs with the highest amounts of Ca-PP (10- and 12.5%), whereas at 52 weeks, the same trend was not observed.

In summary, the present study was designed to determine the impact of a multicomponent CaP material comprising 0-to 12.5% Ca-PP on heterotopic bone formation and the fate of the material *in vivo*. We showed that the presence of Ca-PP does not hinder heterotopic bone formation and has a negligible role in the degradation of the CaP material. However, larger amounts of Ca-PP negatively influenced the level of heterotopic bone mineralisation. The incorporation of Ca-PP within the bone extracellular matrix may impair the mechanical properties of bone that were not explored in this work. Lastly, we confirmed that Ca-PP has low solubility *in vivo*, with phagocytosis being the main mode of removal, although the long-term impact of phagocytosed Ca-PP and the fate of phagocytising multinucleated cells remains to be determined. Phagocytised Ca-PP could accumulate in the surrounding tissues long after the degradation of the implanted material. Notably, Ca-PP is among the main impurities associated with β-TCP synthesis, with the International Organisation for Standardisation (ISO) standard (13175-3:2012) describing any β-TCP product with up to 5 wt% of foreign phase as pure (59). Long-term *in vivo* studies are needed to determine the fate of biomaterials containing Ca-PP, including the fate of degradation products, as well as the influence of Ca-PP on bulk mechanical properties of *de novo* bone.

## 4. Materials and Methods

### 4.1. Experimental setup

The multi-component CaP material was made from monetite-forming calcium-based precursor powder comprising monocalcium phosphate monohydrate [Ca(H_2_PO_4_)_2_•H_2_O, MCPM; Alfa Aesar, Thermo Fisher] and beta-tricalcium phosphate [β-Ca_3_(PO_4_)_2_, β-TCP; Sigma-Aldrich] with the addition of dicalcium pyrophosphate (Ca_2_P_2_O_7_, Ca-PP; Sigma-Aldrich) and glycerol (Sigma-Aldrich) as described previously (31–33). To investigate the influence of Ca-PP on heterotopic bone formation and CaP degradation, five material compositions were made (**Supplementary Table 1**) by keeping MCPM and β-TCP in a consistent ratio and adding an increasing amount of Ca-PP powder (0, 3, 6, 10, and 12.5%). The CaP-glycerol paste was moulded over 3D-printed Ti6Al4V ELI frames (6 ×18 ×18 mm) and allowed to set overnight in sterile water. The constructs were removed from the moulds and placed in sterile water for an additional 48 hours to remove excess glycerol. The final constructs were cut to form six connecting CaP tiles of varying sizes (**Figure 9A**). Control implants were made entirely of 3D-printed Ti6Al4V ELI framed mesh with dimensions comparable to those of the experimental CaP implants. All implants were steam sterilised in an autoclave operated at 121 °C for 20 minutes.

**Figure 9.**
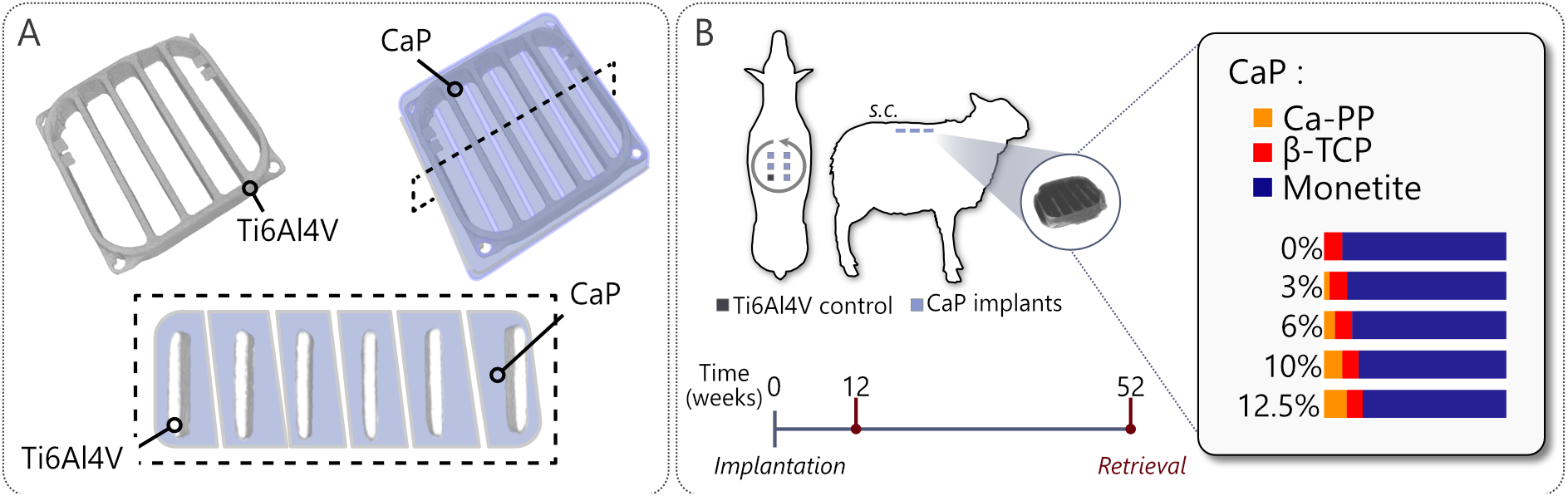
Experimental setup. (A) Calcium phosphate (CaP) implant design with an embedded Ti6Al4V ELI frame. (B) Subcutaneous placement (s.c.) of 5 experimental, CaP, and control (Ti6Al4V only) constructs in 12 adult female sheep (*n* = 6 sheep/time point). Study timeline and CaP composition.

### 4.2. Animal experiment and sample processing

Twelve adult, skeletally mature female sheep (*Ovis aries*) were subcutaneously implanted. The implants were placed at a non-osseous site between the skin and the muscle of the sheep to allow us to investigate the true osteoinductive potential of the CaP material. Anaesthesia in the animals was induced via intravenous propofol injection and maintained with isoflurane inhalation. Under anaesthesia, the selected surgical site was clipped free of wool, cleaned with povidone iodine, and draped to maintain a sterile field. Three incisions were made on each side of the dorsal area, parallel to the vertebral column, to create subcutaneous pockets with blunt dissection. Each sheep received five different CaP material constructs consisting of monetite, β-TCP, with an increasing amount of Ca-PP (0, 3, 6, 10, and 12.5% Ca-PP), as well as a single Ti6Al4V control implant. The exact location of the six implants was decided according to a randomisation scheme (**Figure 9B**). To prevent movement under the skin, the implants were attached to the back muscles of the animals by a surgical thread guided through lateral hoops built into the Ti6Al4V ELI frame. The animals received an intramuscular injection of buprenorphine each day for two days after the surgery.

The anti-inflammatory drug, flunixin, was given for 7 days post-surgery, and antibiotics, amoxicillin and enrofloxacin, for 6 weeks post-surgery. After 12- and 52 weeks (*n* = 6 sheep / time point), the animals were sacrificed via pentobarbital intravenous injection. Implants were dissected *en bloc* with the surrounding soft tissue, fixed in 10% neutral buffered formalin, dehydrated in ethanol, and resin embedded (LR White, London Resin Co. Ltd., UK). The animal experiments were approved by the Ministry of National Education, Higher Education and Research (NAMSA, Chasse-sur-Rhône, France; Approval nr. 01139.2).

### 4.3. Histology and histomorphometry

Undecalcified, toluidine blue and Van Gieson-stained histological sections (∼40 µm thick) were prepared from bisected, resin-embedded blocks. Areas where collagen is present stain positively with Van Gieson’s stain, and therefore, collagenous sites (i.e., heterotopic bone) and non-collagenous sites (i.e., CaP) can be readily distinguished. Toluidine blue stain was used in a second set of histological sections to determine the presence and involvement of different cellular components. Brightfield imaging was performed using a Nikon Eclipse E600 optical microscope (Nikon Ltd., Tokyo, Japan). Van Gieson-stained sections were imaged using 10× magnification and used to measure the heterotopic B.Ar. and CaP.Ar. present in the sections. Area measurements were made in ImageJ (imagej.nih.gov/ij) by manually demarcating the areas of interest – heterotopic bone and the CaP material.

### 4.4. Raman spectroscopy

The remaining halves of the resin-embedded blocks were wet polished using 400–4000 grit silicon carbide (SiC) paper, with the final polishing step performed with absolute ethanol. Initial identification of ROIs was made using BSE-SEM in a Quanta 200 environmental SEM (FEI Company, The Netherlands) operated at 20 kV acceleration voltage, 1 Torr water vapour pressure, and 10 mm working distance (**Supplementary Figures 1-10**). Three distinct ROIs were selected: (i) heterotopically formed bone, (ii) CaP located 50– 100 µm from the bone interface, and (iii) CaP areas farthest from the bone interface, typically within 50 µm from the Ti6Al4V ELI frame. Additionally, energy dispersive X-ray spectroscopy (EDX) maps of phagocytosed CaP material (264 × 214 µm) were made in the inter-tile area of two representative samples, one for each time point, at 15kV, 1 Torr water vapour pressure, and 10 mm working distance. For chemical composition analysis of the CaP material and heterotopic bone, Raman spectroscopy was performed using a confocal Raman microscope (Renishaw inVia Qontor, Renishaw plc. Wotton under Edge, UK) equipped with a 785 nm laser and Live-Track focus-tracking technology for enhanced signal stability (34). The laser was focused on the sample surface through a 100x/0.9 NA objective, and the Raman scattered light was collected using a Peltier-cooled charge-coupled device deep depletion near-infrared enhanced detector behind a 1200 g mm^−1^ grating. The laser power at the sample was ∼10 mW. On each sample (*n* = 6), six point measurements were obtained at a 2 s integration time and 5 accumulations in each ROI. Background subtraction and cosmic ray removal were performed in Spectragryph V1.2.16.1 (**Supplementary Figure 11**) (35). In heterotopic bone, mineral crystallinity corresponds to the inverse value of full-width at half-maximum (1/FWHM) of 𝓋_1_PO_4_^3-^ peak (960 cm-1). The carbonate-to-phosphate ratio is the intensity ratio between the 𝓋_1_CO_3_^2-^ (∼1070 cm-1) and 𝓋_1_PO_4_^3-^ (∼960 cm^−1^) peaks. The mineral-to-matrix ratio is the integral area ratio between the 𝓋_2_PO_4_^3-^ (420–470 cm-^1^) peak and the amide III band (1240–1270 cm^−1^).

### 4.5. Statistical analysis

Statistical analyses were performed using GraphPad Prism v10.0.0 (GraphPad Software, USA). The non-parametric Mann-Whitney U test was used for comparisons between groups, and *p* values <0.05 were considered statistically significant. Mean values ± standard deviations (SD) are presented. Spearman correlation analysis was performed between the B.Ar. and CaP.Ar. Based on the value of the Spearman’s rank coefficient, the strength of the correlation is interpreted as: little if any correlation (0.00–0.30), low (0.30–0.50), high (0.70–0.90), and very high (0.9–1.00).

## Acknowledgements

The authors would like to thank Marc Bohner for scientific input and critical reading of this manuscript. Lena Emanuelsson and Birgita Norlindh for histological sample preparation. This work was financially supported by the Svenska Sällskapet för Medicinsk Forskning (SSMF), the Swedish Research Council (2017-04728, 2018-02891, and 2020-04715), the Swedish Foundation for Strategic Research (RMA15-0110), the Swedish state under the agreement between the Swedish government and the county councils, the ALF agreement (ALFGBG-725641), the IngaBritt and Arne Lundberg Foundation, the Hjalmar Svensson Foundation, the Adlerbertska Foundation, the Kungliga Vetenskaps-och Vitterhets-Samhället i Göteborg, the Eivind och Elsa K:son Sylvans Stiftelse, and the Area of Advance Materials of Chalmers and GU Biomaterials within the Strategic Research Area initiative launched by the Swedish government. The funding sources had no role in the conceptualisation, design, data collection, analysis, decision to publish, or preparation of the manuscript.

## Data availability

The data will be available upon reasonable request.

## Author contributions

Martina Jolic: Methodology, Investigation, Visualization, Writing – original draft, Funding acquisition. Isabella Åberg: Methodology. Omar Omar: Conceptualization, Animal surgery. Håkan Engqvist: Conceptualization, Funding acquisition. Thomas Engstrand: Conceptualization. Anders Palmquist: Investigation, Supervision, Writing – original draft, Funding acquisition. Peter Thomsen: Conceptualization, Supervision, Writing – original draft, Funding acquisition. Furqan A. Shah: Conceptualization, Supervision, Writing – original draft, Funding acquisition.

