## Supplementary Document for "Pyrophosphate-containing Calcium Phosphates Negatively Impact Heterotopic Bone Quality"

**Supplementary Table 1.** *Implanted CaP material formulations.*

| Group | Components (in %) | | |
|---|---|---|---|
| | Monetite | β-TCP | Ca-PP |
| 0% Ca-PP | 90.9 | 9.1 | 0 |
| 3% Ca-PP | 88.18 | 8.81 | 3 |
| 6% Ca-PP | 85.45 | 8.54 | 6 |
| 10% Ca-PP | 81.82 | 8.18 | 10 |
| 12.5% Ca-PP | 79.55 | 7.95 | 12.5 |

**Supplementary Figure 1.** *Partial overviews of undecalcified, bisected sample blocks imaged with backscattered electron scanning electron microscopy (BSE-SEM) used to identify regions of interest for Raman spectroscopy. Samples of 0% Ca-PP constructs at 12 weeks. Scale bars = 1 mm.*

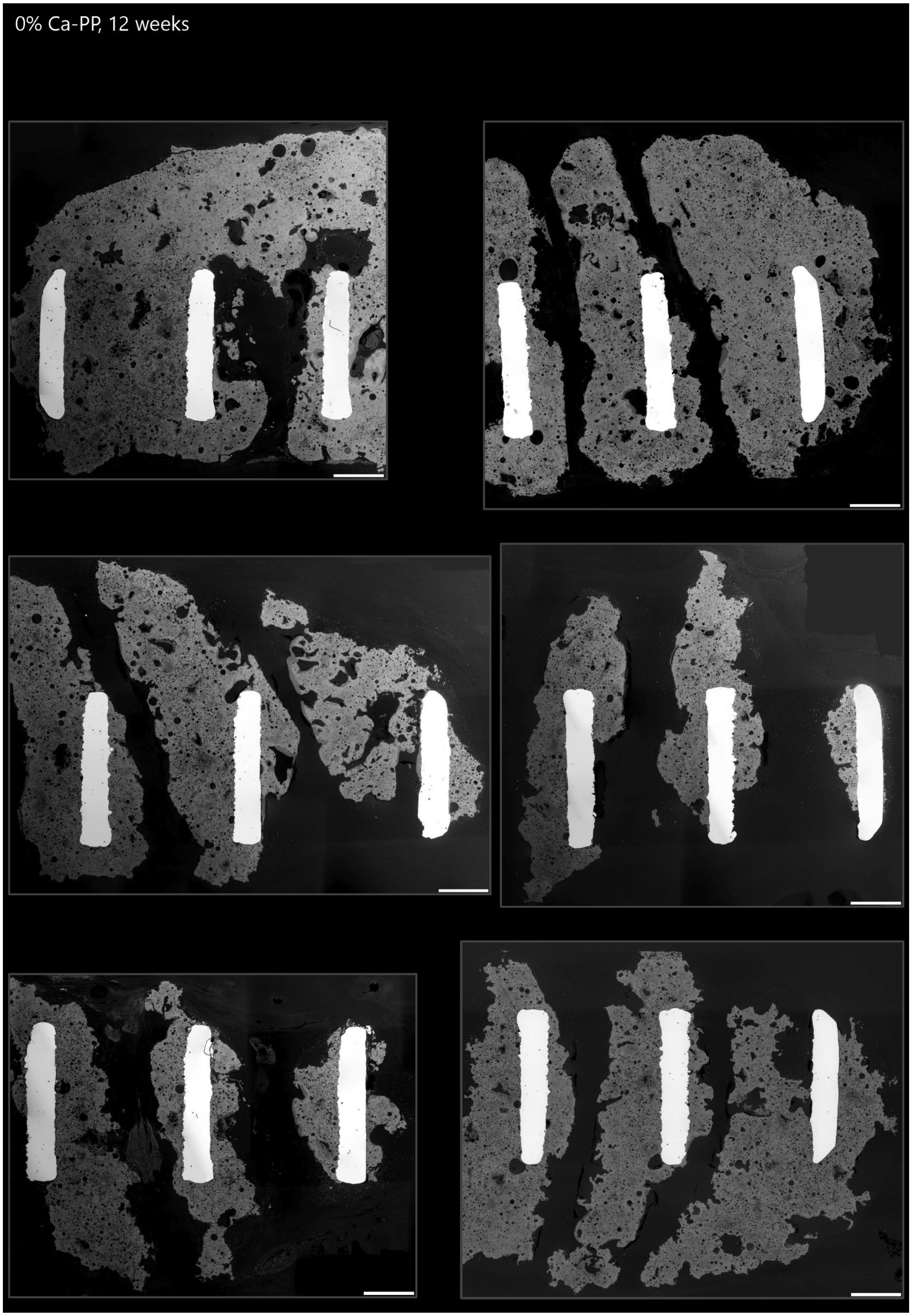

**Supplementary Figure 2.** *Partial overviews of undecalcified, bisected sample blocks imaged with BSE-SEM used to identify regions of interest for Raman spectroscopy. Samples of 3% Ca-PP constructs at 12 weeks. Scale bars = 1 mm.*

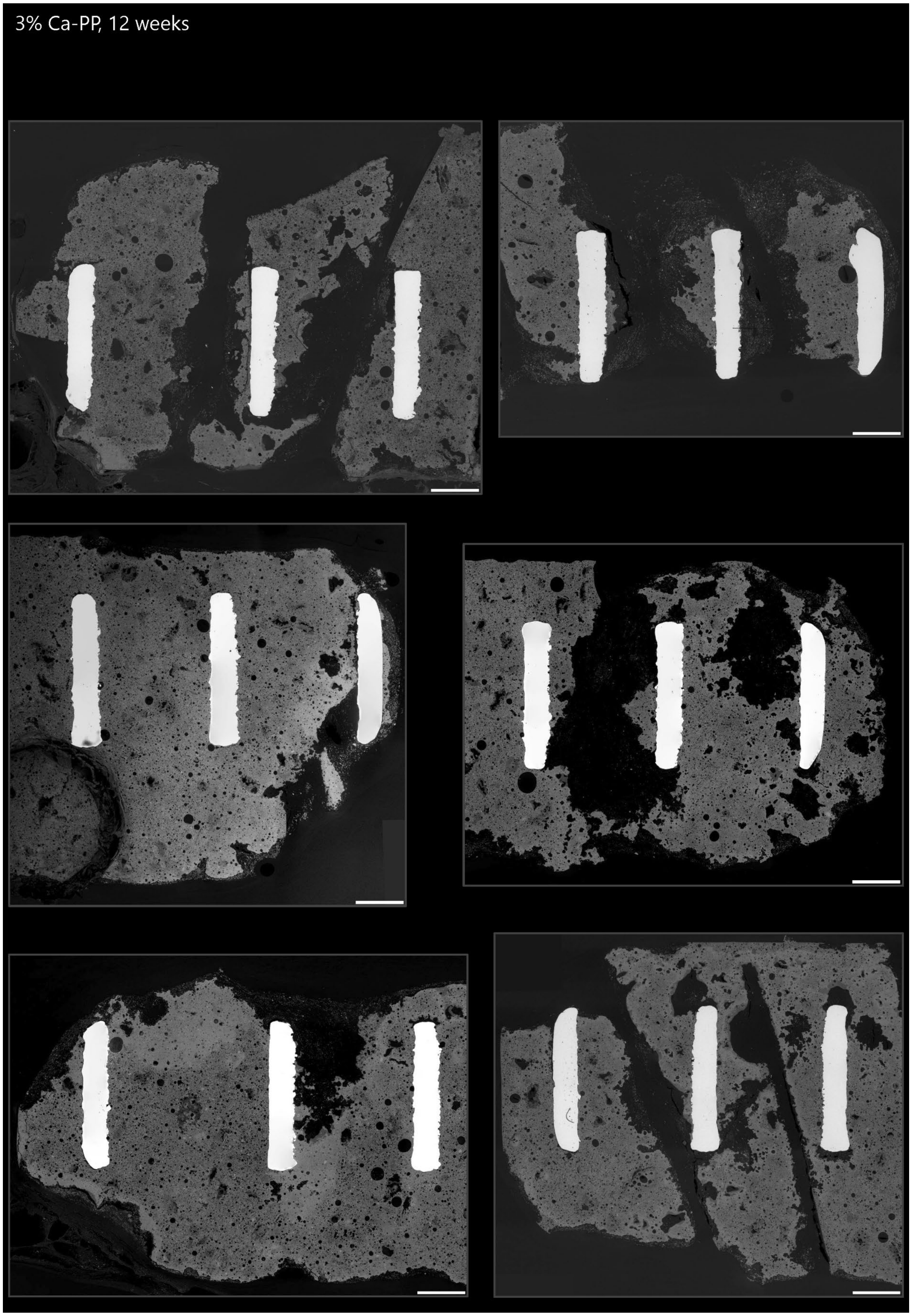

**Supplementary Figure 3.** *Partial overviews of undecalcified, bisected sample blocks imaged with BSE-SEM used to identify regions of interest for Raman spectroscopy. Samples of 6% Ca-PP constructs at 12 weeks. Scale bars = 1 mm.*

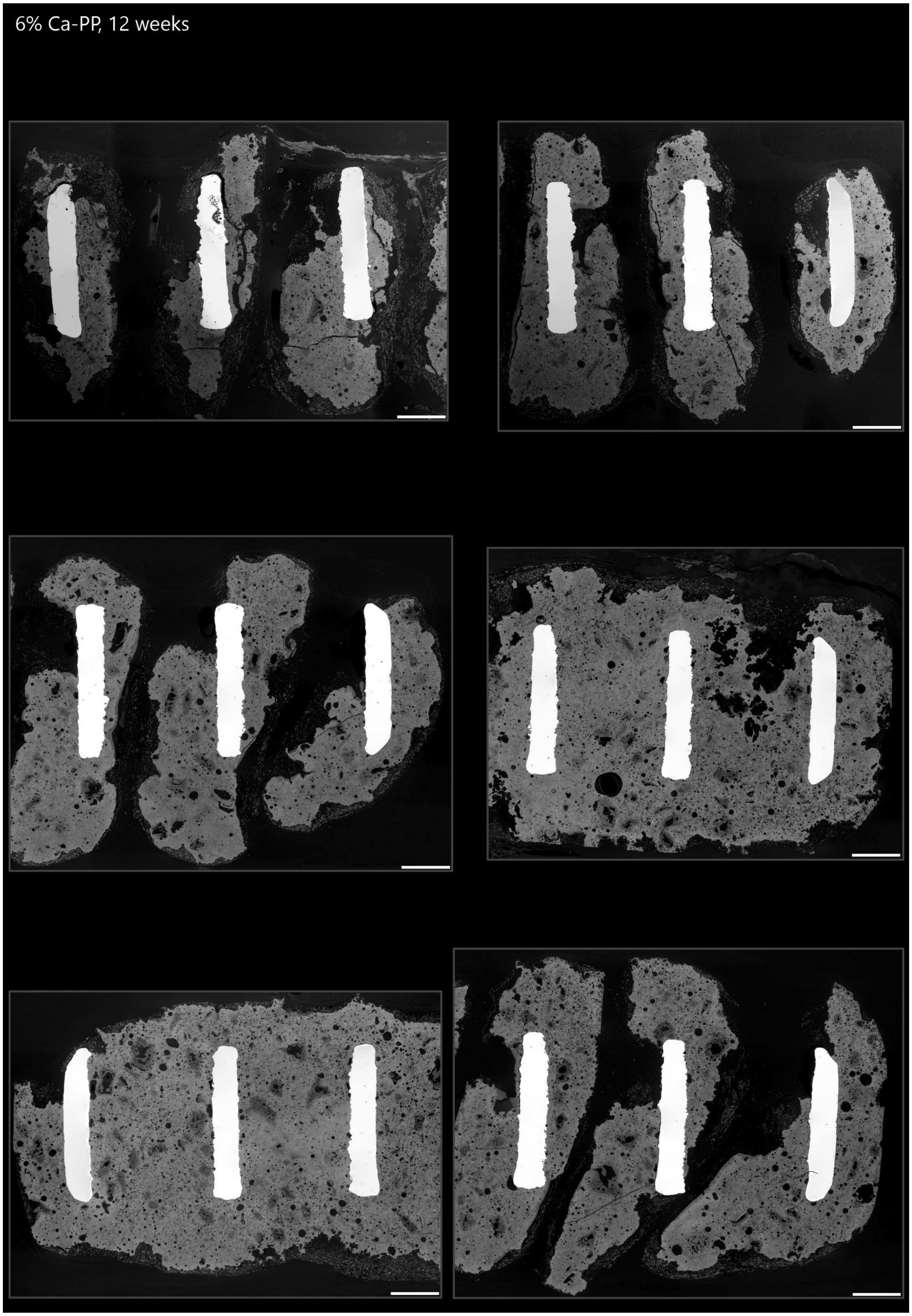

**Supplementary Figure 4.** *Partial overviews of undecalcified, bisected sample blocks imaged with BSE-SEM used to identify regions of interest for Raman spectroscopy. Samples of 10% Ca-PP constructs at 12 weeks. Scale bars = 1 mm.*

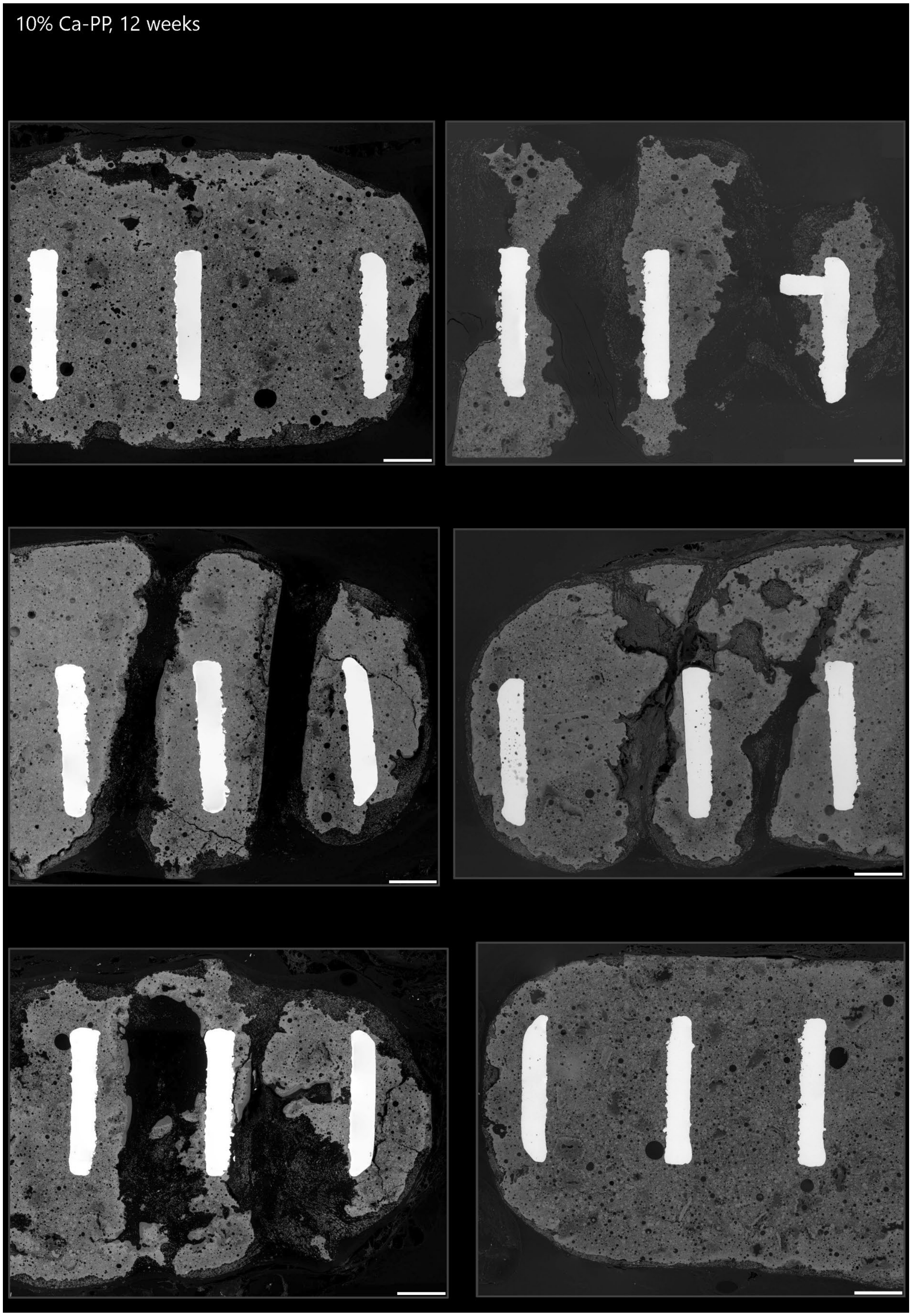

**Supplementary Figure 5.** *Partial overviews of undecalcified, bisected sample blocks imaged with BSE-SEM used to identify regions of interest for Raman spectroscopy. Samples of 12.5% Ca-PP constructs at 12 weeks. Scale bars = 1 mm.*

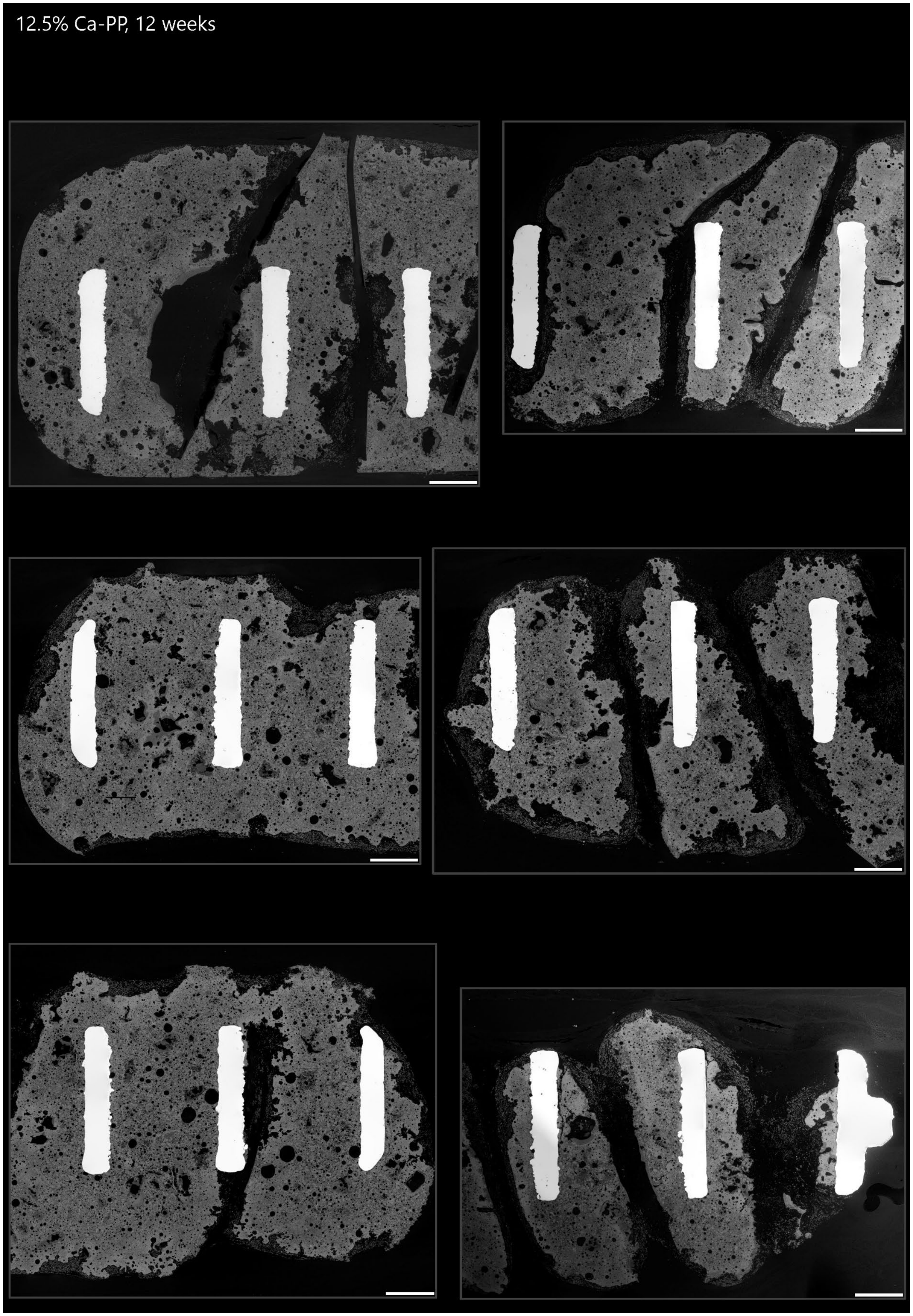

**Supplementary Figure 6.** *Partial overviews of undecalcified, bisected sample blocks imaged with BSE-SEM used to identify regions of interest for Raman spectroscopy. Samples of 0% Ca-PP constructs at 52 weeks. Scale bars = 1 mm.*

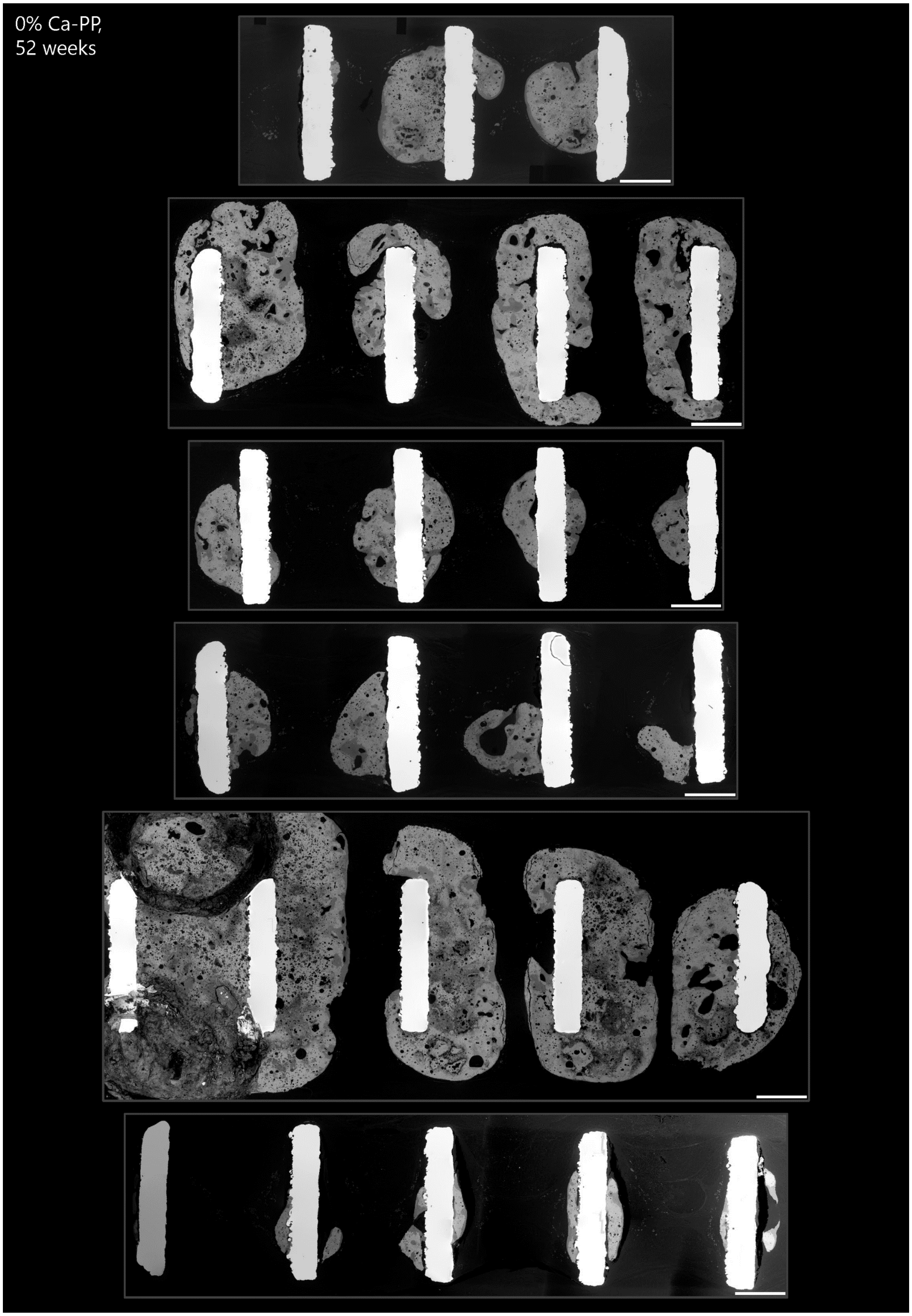

**Supplementary Figure 7.** *Partial overviews of undecalcified, bisected sample blocks imaged with BSE-SEM used to identify regions of interest for Raman spectroscopy. Samples of 3% Ca-PP constructs at 52 weeks. Scale bars = 1 mm.*

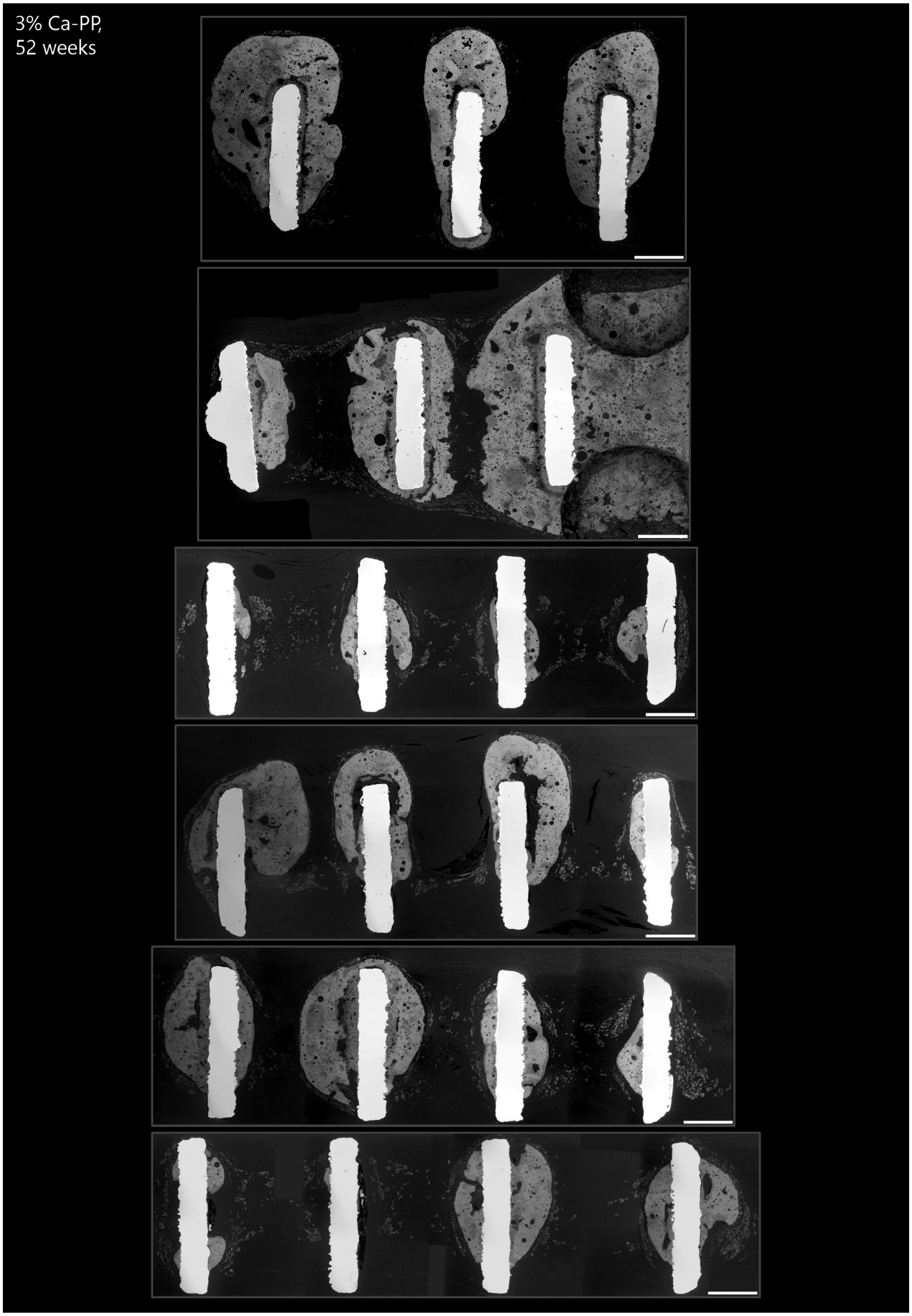

**Supplementary Figure 8.** *Partial overviews of undecalcified, bisected sample blocks imaged with BSE-SEM used to identify regions of interest for Raman spectroscopy. Samples of 6% Ca-PP constructs at 52 weeks. Scale bars = 1 mm.*

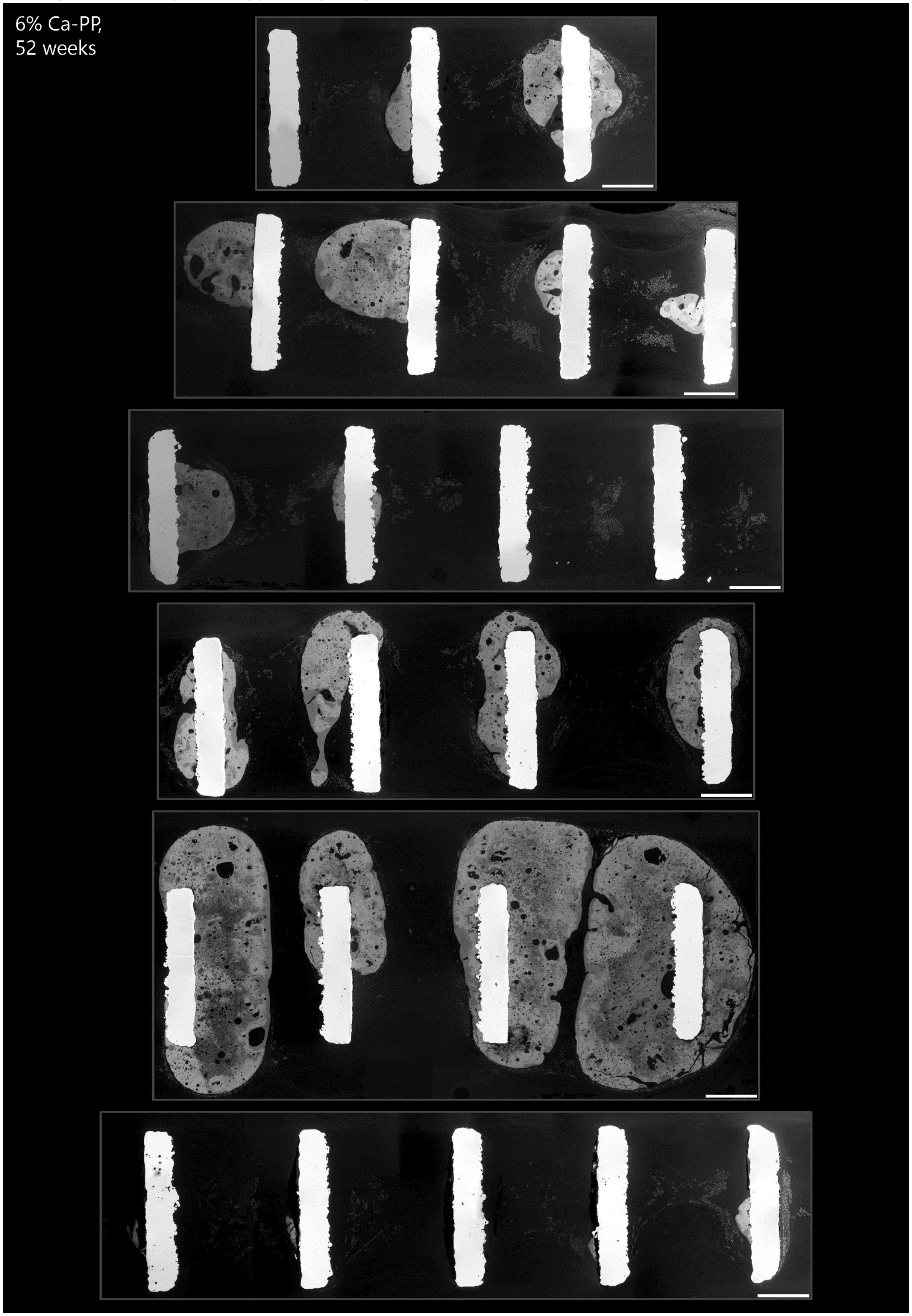

**Supplementary Figure 9.** *Partial overviews of undecalcified, bisected sample blocks imaged with BSE-SEM used to identify regions of interest for Raman spectroscopy. Samples of 10% Ca-PP constructs at 52 weeks. Scale bars = 1 mm.*

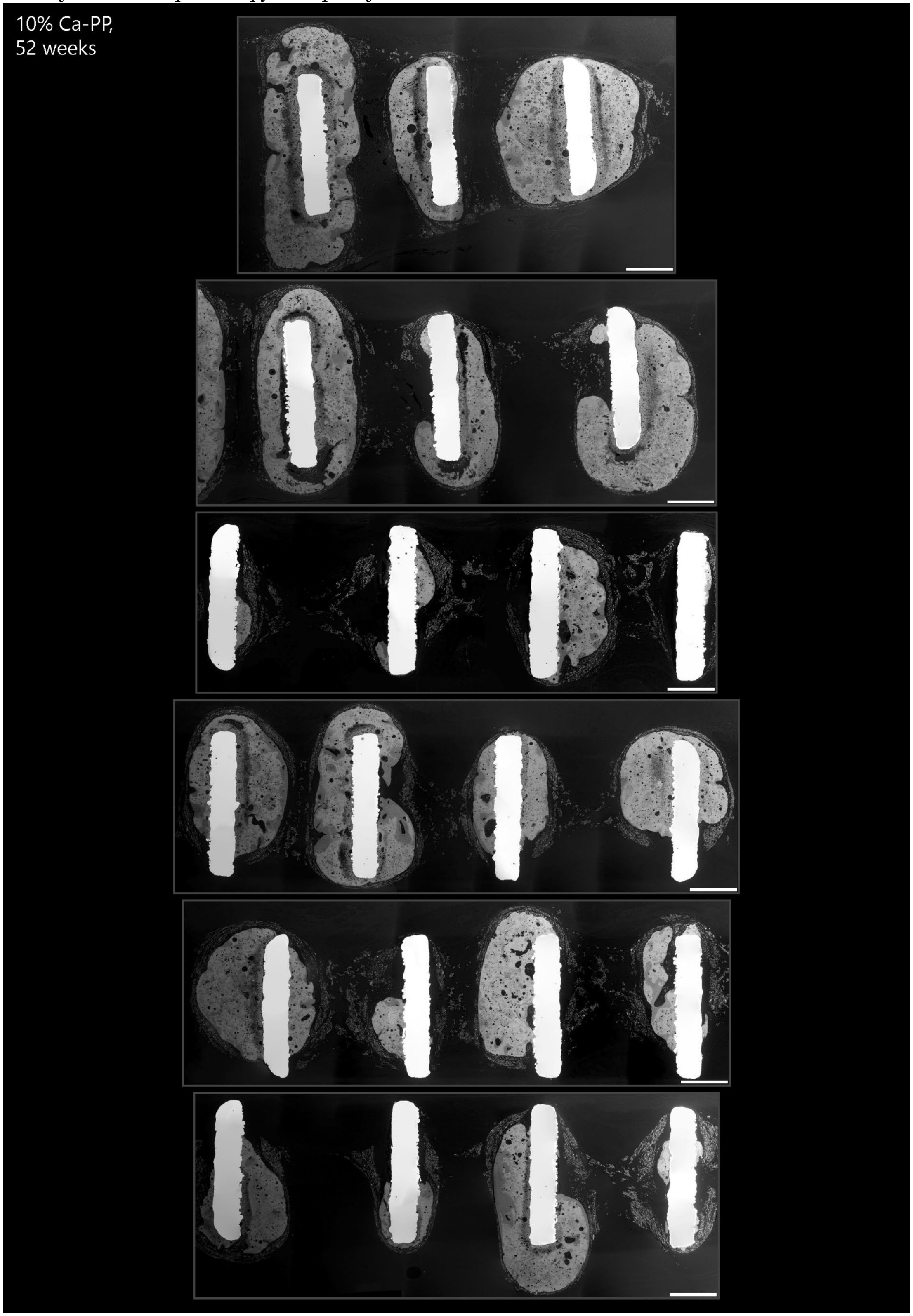

**Supplementary Figure 10.** *Partial overviews of undecalcified, bisected sample blocks imaged with BSE-SEM used to identify regions of interest for Raman spectroscopy. Samples of 12.5% Ca-PP constructs at 52 weeks. Scale bars = 1 mm.*

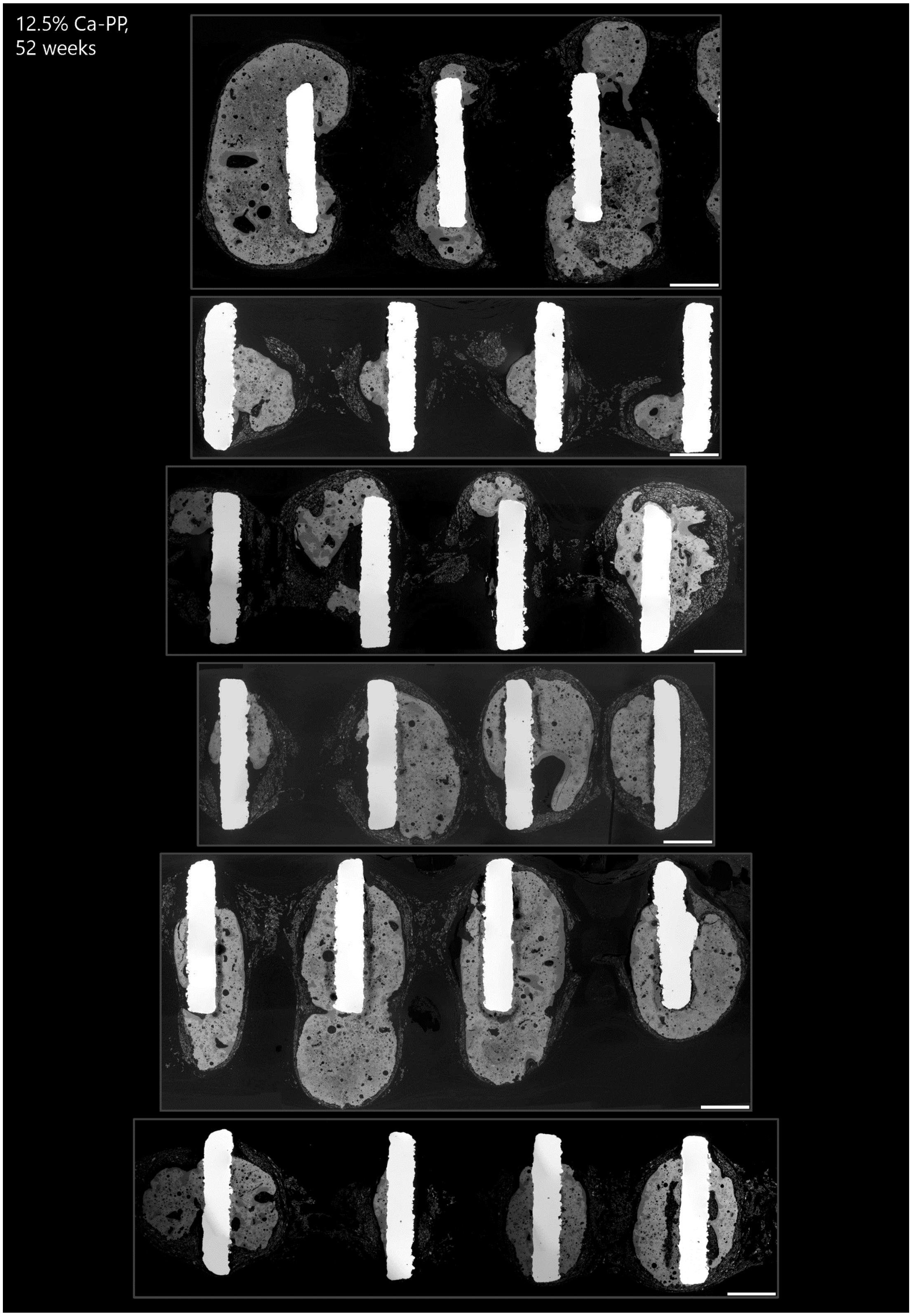

**Supplementary Figure 11.** *Representative Raman spectra obtained from the three regions of interest: heterotopic bone, CaP at the bone interface, and CaP furthest away from the bone interface at 12- and 52 weeks in vivo. Acquired Raman spectra were processed by performing background subtraction at 20% coarseness and cosmic ray removal with remove spikes function at maximum width at 5 pixels in Spectragryph V1.2.16.1. Raw spectrum (green), background fluorescence profile (red), and corrected spectrum (blue).*

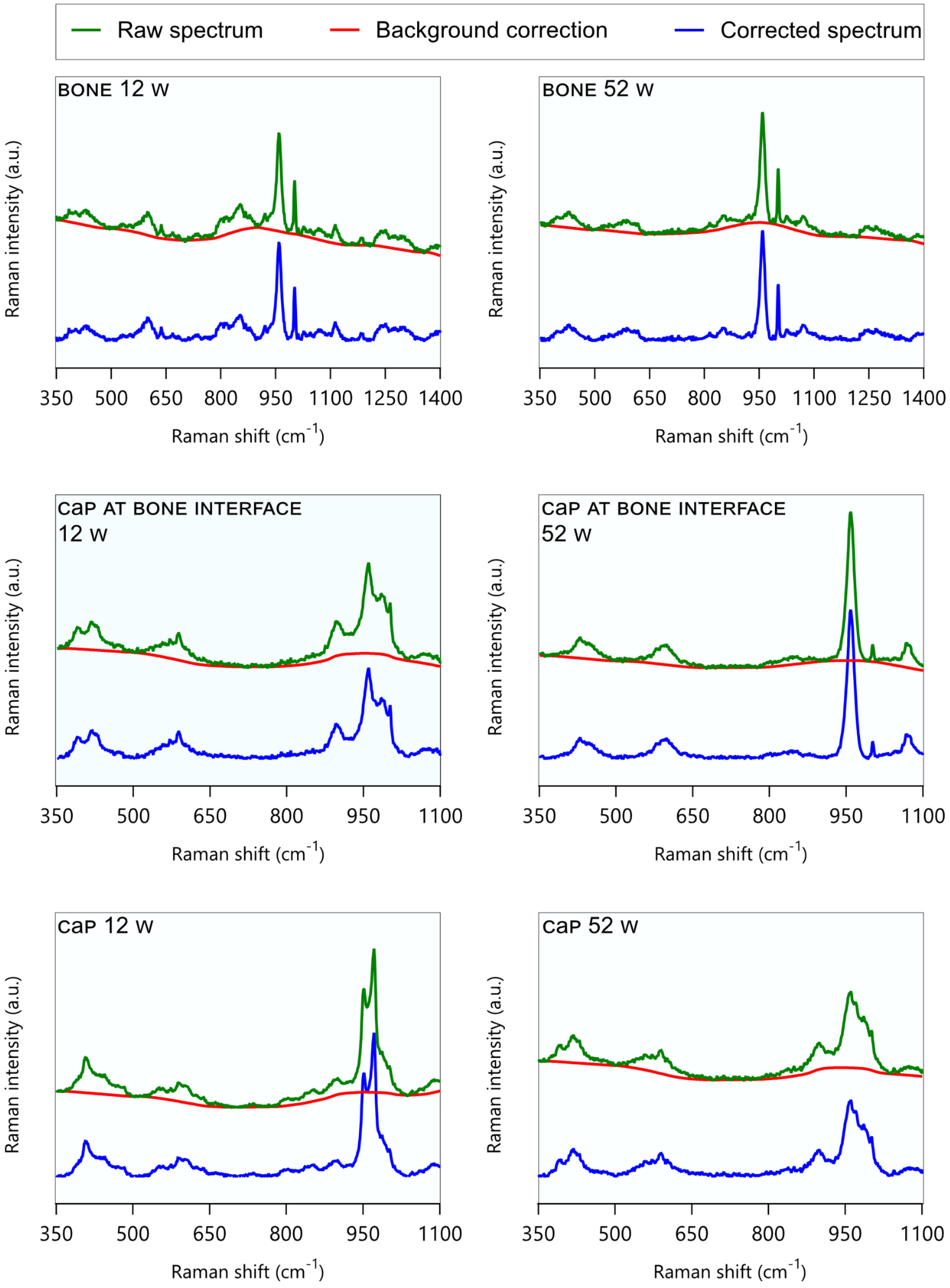

**Supplementary Figure 12.** *Overviews of undecalcified, histological sections of control Ti6Al4V ELI implants stained with Van Gieson at 12 weeks (n = 6). Scale bars = 1 mm.*

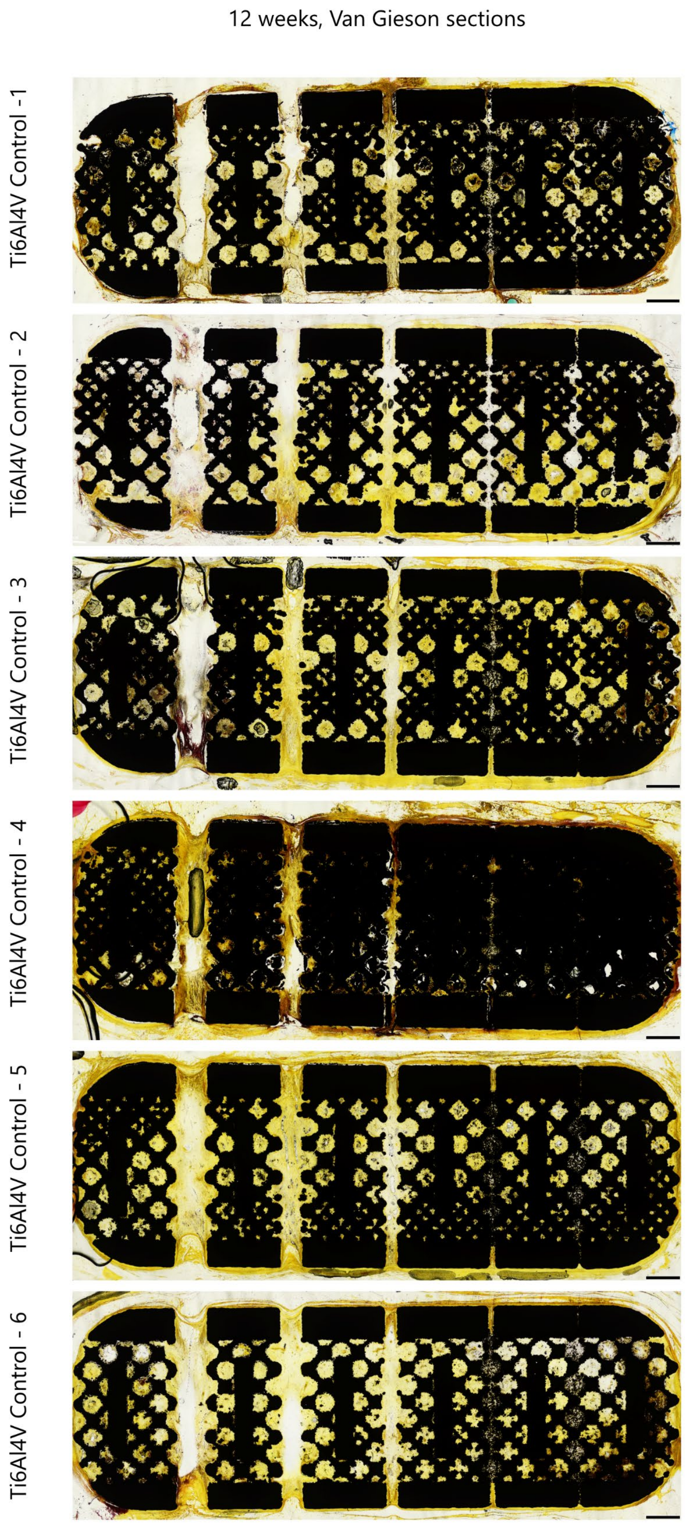

**Supplementary Figure 13.** *Overviews of undecalcified, histological sections of control Ti6Al4V ELI implants stained with Van Gieson at 52 weeks (n = 6). Scale bars = 1 mm.*

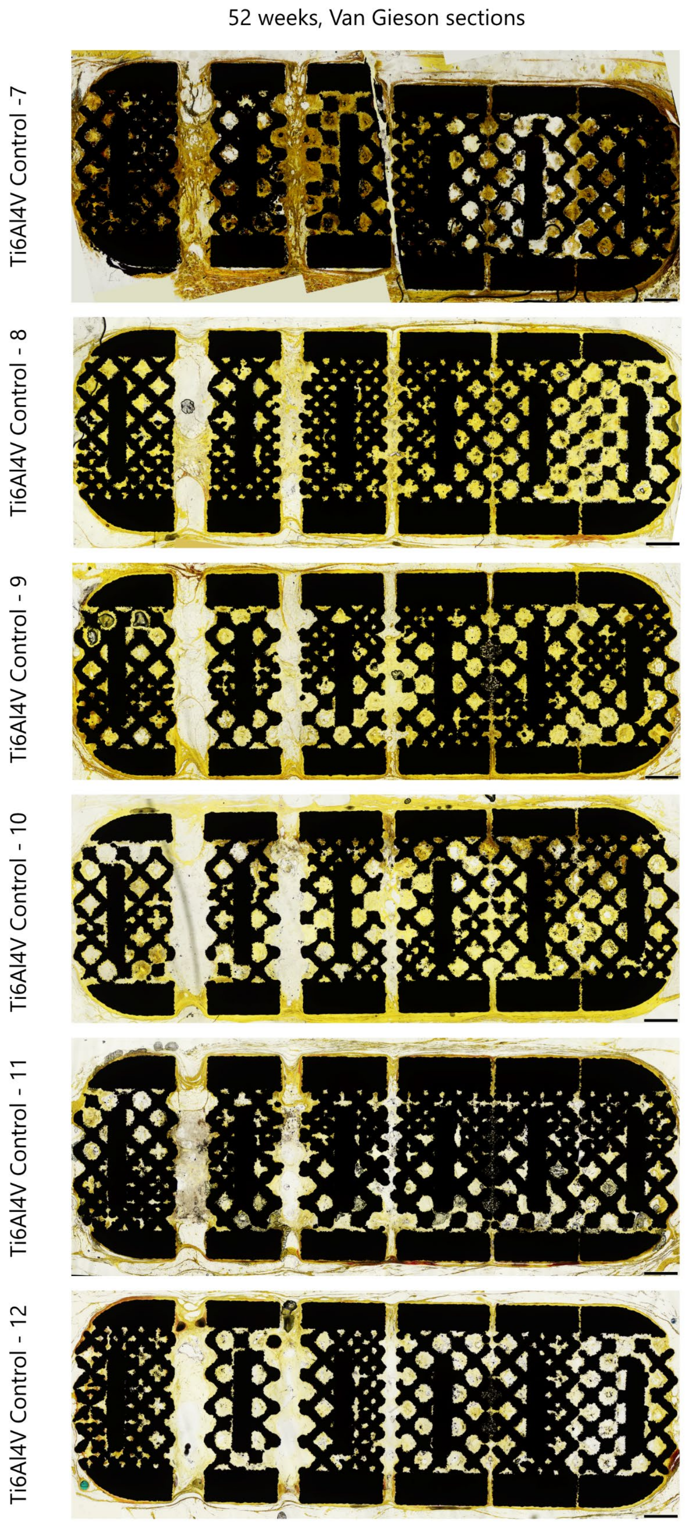

**Supplementary Table 2.** *Measured bone areas (B.Ar.).*

| B.Ar. (mm$^2$) | *12 weeks in vivo* | | | | |
|---|---|---|---|---|---|
| Animal # | 0% Ca-PP | 3% Ca-PP | 6% Ca-PP | 10% Ca-PP | 12.5% Ca-PP |
| 1 | 1.12 | 0.23 | 0.78 | 2.43 | 0.51 |
| 2 | 0.40 | 0.15 | 0.32 | 0.05 | 0.19 |
| 3 | 0.38 | 0.07 | 0.18 | 0.29 | 0.45 |
| 4 | 0.37 | 1.55 | 0.99 | 0.99 | 0.58 |
| 5 | 2.64 | 1.10 | 1.29 | 2.96 | 2.04 |
| 6 | 0.91 | 0.35 | 0.78 | 1.13 | 0.67 |
| | *52 weeks in vivo* | | | | |
| 7 | 1.43 | 0.55 | 1.01 | 1.73 | 0.55 |
| 8 | 0.44 | 3.80 | 0.22 | 1.65 | 1.26 |
| 9 | 3.04 | 2.09 | 0.79 | 2.77 | 2.19 |
| 10 | 3.18 | 2.45 | 2.45 | 2.75 | 5.37 |
| 11 | 1.15 | 0.75 | 0.39 | 1.40 | 1.21 |
| 12 | 0.74 | 1.24 | 0.12 | 1.17 | 0.95 |

**Supplementary Table 3.** *Measured calcium phosphate areas (CaP.Ar).*

| CaP.Ar. (mm$^2$) | *12 weeks in vivo* | | | | |
|---|---|---|---|---|---|
| Animal # | 0% Ca-PP | 3% Ca-PP | 6% Ca-PP | 10% Ca-PP | 12.5% Ca-PP |
| 1 | 61.83 | 72.77 | 50.06 | 50.21 | 63.21 |
| 2 | 36.60 | 24.83 | 35.20 | 45.33 | 49.26 |
| 3 | 38.72 | 48.59 | 16.02 | 51.32 | 59.13 |
| 4 | 48.01 | 59.03 | 36.95 | 53.68 | 46.26 |
| 5 | 54.32 | 70.42 | 47.38 | 53.62 | 49.15 |
| 6 | 26.18 | 27.46 | 38.89 | 51.97 | 46.18 |
| | *52 weeks in vivo* | | | | |
| 7 | 6.72 | 1.66 | 5.13 | 11.00 | 2.93 |
| 8 | 4.21 | 35.79 | 1.60 | 10.01 | 7.89 |
| 9 | 17.55 | 25.79 | 7.50 | 27.51 | 20.67 |
| 10 | 35.12 | 24.35 | 41.62 | 26.01 | 25.13 |
| 11 | 5.37 | 3.23 | 3.05 | 7.92 | 4.83 |
| 12 | 1.56 | 7.36 | 0.54 | 3.03 | 7.39 |

**Supplementary Figure 14.** *Energy-dispersive X-ray spectroscopy (EDX) maps of representative regions of interest of phagocytised CaP debris at 12- and 52 weeks in vivo. Elemental maps for Ca, C, P, and merged confirm that phagocytied debris is indeed CaP. Insets: summary spectra of the EDX map. Scale bars = 100 μm.*

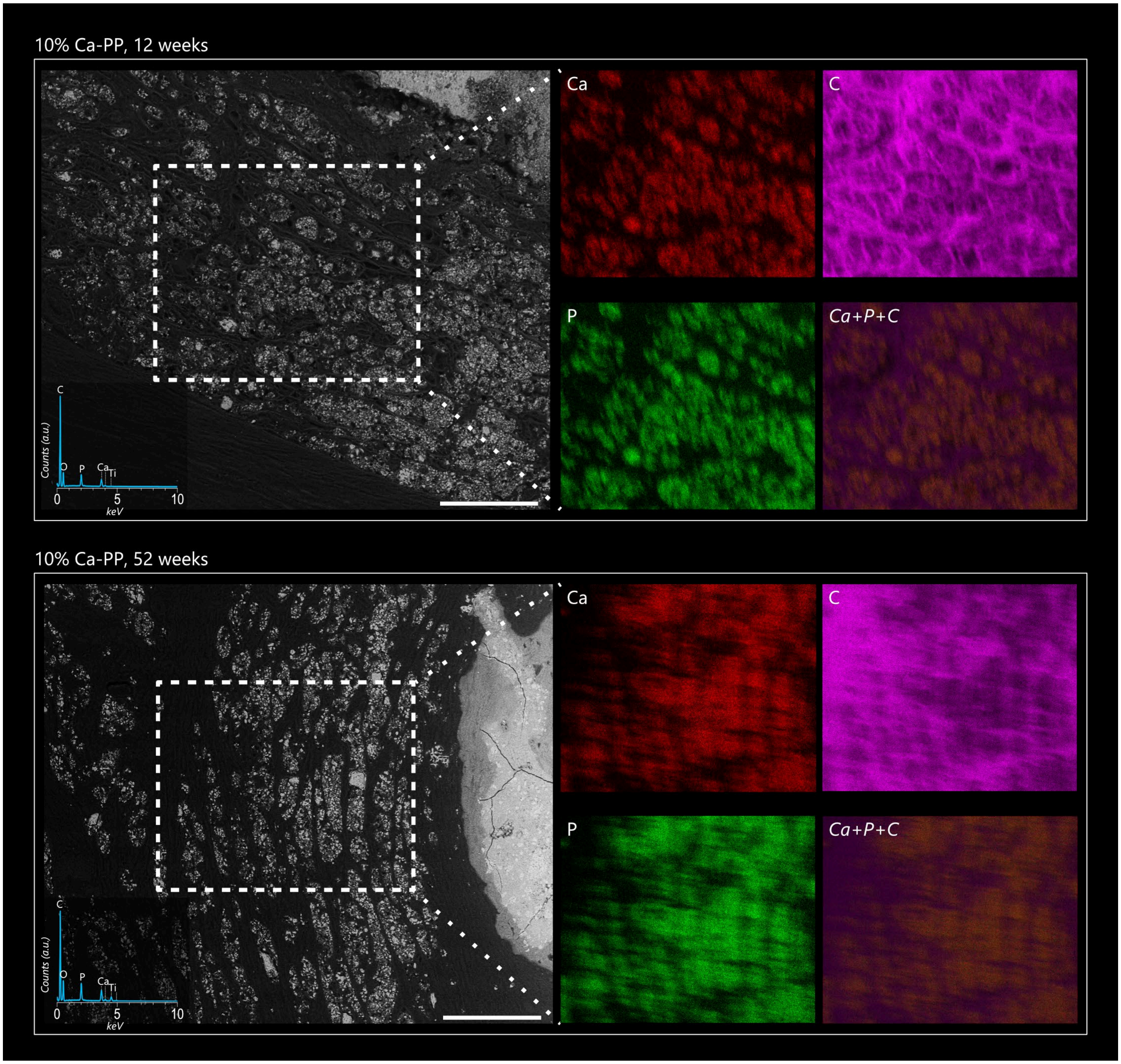
